# Prediction of LncRNA Encoded Small Peptides in Glioma and The Oligomer Channel Functional Analysis Using in Silico Approaches

**DOI:** 10.1101/2020.05.13.094763

**Authors:** Yipeng Cao, Rui Yang, Imshik Lee, Wenwen Zhang, Jiana Sun, Wei Wang

**Affiliations:** Tianjin Medical University Cancer Institute and Hospital, National Clinical Research Center for Cancer, Tianjin, 300060 P.R. China; College of Physics, Nankai University, Tianjin, 300071 P.R. China; National Supercomputer Center in Tianjin, 300457 P.R. China; Tianjin Union Medical Center, Nankai University Affiliated Hospital. 190, Jieyuan Road, Hongqiao District, Tianjin, 300031, P.R. China

**Keywords:** MD simulation, ion channel, lncRNA, ceRNA, glioma, small peptides

## Abstract

Glioma is lethal malignant brain cancers, many reports have shown that abnormalities in the behavior of water and ion channels play an important role in regulating glioma proliferation, migration, apoptosis and differentiation. Recently, new studies have suggested that some long noncoding RNAs (lncRNAs) containing small open reading frames (smORFs), can encode small peptides and form oligomers for water or ion regulation. However, because these peptides are difficult to identify, their functional mechanisms are far from being clearly understood. In this study, we used bioinformatic methods and softwares to identify and evaluate lncRNAs in gliomas that may encode small transmembrane peptides. Combining ab initio homology modeling, molecular dynamics simulations and energetic calculations, we constructed a predictive model and predicted the oligomer channel activity of peptides by identifying the lncRNA ORFs.

We found that one key hub lncRNA, DLEU1, which contains two smORFs (ORF1 and ORF8) could encode small peptides that form pentameric channels. The mechanics of water and ion (Na + and Cl-) transport through this pentameric channel were simulated. The potential of mean force (PMF) of the H_2_O molecules along the two ORF-encoded peptide channels indicated that the energy barrier was different between ORF1 and ORF8. The ORF1-encoded peptide pentamer acted as a self-assembled water channel but not as an ion channel, and the ORF8 neither permeating ions nor water.

This work provides new methods and theoretical support for further elucidation of the function of lncRNA-encoded small peptides and their role in cancer. Additionally, it provides a theoretical basis for drug development.

## 1, Introduction

Glioma is one of the most prevalent types of primary intracranial carcinomas; it has varying malignancy grades and histological subtypes ^1^ and remains a highly lethal malignancy worldwide. Glioma is characterized by rapid cell proliferation and angiogenesis ^2^. Traditional treatments have limited effectiveness in the majority of gliomas because glioblastoma (GBM) stem-like cells (GSCs) are highly recrudescent ^3^. Consequently, investigations exploring the accurate molecular mechanisms of and reliable therapeutic targets for gliomas have attracted extensive attention.

Many studies show that long noncoding RNAs (lncRNAs) are involved in many diseases, including glioma, and participate in gene regulation via various mechanisms ^4 5^. LncRNAs are a class of RNAs > 200 nucleotides (nt) in length that lack coding potential ^6 7^. Based on the increasing amount of functional lncRNAs aberrantly expressed in glioma tissues and cell lines ^8 9 10^, they closely related to glioma occurrence by influencing tumor development, invasion, and metastasis. Recently, some studies reported the discovery of ORFs that encode functional small peptides in particular lncRNAs ^11 12^; for example, Huang et al. found that the lncRNA HOXB-AS3 encodes micropeptides that can bind to heterogeneous nuclear ribonucleoproteins (hnRNPs) and suppress glucose metabolism reprogramming in colon cancer ^13^. SLN, PLB, MLN, and DWORF ^14 15^ are involved in some physiological processes of diseases by regulating skeletal muscle activity to interact with sarcoplasmic reticulum Ca^2+^-ATPase (SERCA) ^16^. Ji et.al surprisingly found that more than 40% of lncRNA fragments encode proteins and that more than half of these fragments are in the form of small peptides (median number of amino acids (AAs) <46) ^17^.

Previous reports showed that glioma proliferation, apoptosis, migration and invasion are driven by water permeability mostly through aquaporins (AQPs) ^18 19^; this information has often been used as a marker of expected survival in cancer patients. Water regulation is strongly correlated with glioma angiogenesis, brain edema and tumor migration and invasion ^20 21^, indicating that study of the water permeability mechanism could lead to the discovery of pharmaceutical targets for glioma management. In addition to AQP water channels, and more importantly, lncRNA-encoded small peptides, such as SLN and PLB, can form pentameric channels in the membrane through the leucine/isoleucine zipper in their transmembrane (TM) regions ^22^. The pentameric channels formed by these peptides act as TM water and ion transport channels and hydrophobic gate channels.

Some reports suggest that the AQPs in gliomas are not permeable enough to cause glioma progression ^23 24 25^. Therefore, we considered that the lncRNA-encoded small peptide oligomers may be involved in water penetration. Because experimental identification of lncRNA-encoded small peptides is also very difficult, the present bioinformatic methods were implemented mainly by algorithms such as support vector machine (SVM) algorithms, depending on the length of the amino acid chain and the homology to known sequences ^12^. However, the encoded small TM peptides contain so few amino acids (=<46 AA) that most are not homologous with other proteins and thus cannot be identified accurately by the current algorithms.

In this study, we identified the key hub lncRNAs in gliomas by multiple bioinformatic methods, including survival analysis, lncRNA functional enrichment analysis, bioinformatic predictions, and ab initio modeling, to improve the predictive reliability of lncRNA-encoded small peptides. Subsequent structural analysis evaluated the possible encoding of small TM peptides. Molecular dynamics (MD) simulation was applied to investigate whether the small TM peptides could stabilize the formation of a pentameric structure acting as a water or ion channel. Our results showed that the peptides encoded by a hub lncRNA formed a pentamer with water and ion permeability, supplementing our understanding of the permeability mechanism of water in gliomas. Thus, our results suggest that the hub lncRNAs in glioma could encode small TM peptides and that their self-assembled oligomers could be functionalized as water channels. Additionally, our study provides new methods and theoretical support for further elucidation of the functions of lncRNA-encoded small peptides and their role in cancer and also provides a theoretical basis for drug development.

## 2. Materials and Methods

### 2.1 Screening and analysis of glioma hub lncRNAs

#### 1) Computational analysis of RNA sequences

All data were downloaded from The Cancer Genome Atlas (TCGA) database (https://portal.gdc.cancer.gov) with complete lncRNA, mRNA and miRNA expression profiles. The total glioma RNA sequences of 700 tumor samples and 5 adjacent nontumorous brain tissues were included in subsequent analysis. The patients had an overall survival time of >10 years. This study followed the TCGA Research Network guidelines (http://cancergenome.nih.gov/publications/publicationguidelines). Thus, no further ethical approval was required.

#### 2) Statistical analysis

Differential expression analysis was performed to identify differentially expressed lncRNAs, mRNAs and miRNAs (DElncRNAs, DEmRNAs, and DEmiRNAs, respectively) by the edgeR package in Bioconductor with the cutoff criteria of |log2 fold change (FC)|>2 and false discovery rate (FDR) <0.01. The *Homo sapiens* GTF file for gene annotation was downloaded from the National Center for Biotechnology Information (NCBI) website (http://ftp.ncbi.nlm.nih.gov/genomes/Homo_sapiens/).

#### 3) Gene Ontology (GO), pathway analysis, and ceRNA network construction

Significant DElncRNAs, DEmRNAs, and DEmiRNAs were examined in the GO database (http://www.geneontology.org), where significantly enriched GO terms were identified to analyze their biological function. A ceRNA network was built including lncRNA-miRNA-mRNA interactions. Human lncRNA-miRNA and mRNA-miRNA interactions were obtained from StarBase 2.0 (http://starbase.sysu.edu.cn/starbase2/index.php). According to the lncRNA-miRNA and miRNA-mRNA interactions, we utilized Cytoscape (version 3.7.0) ^26^ to visualize the lncRNA-miRNA-mRNA network.

#### 4) Survival differences of hub lncRNAs

From the obtained ceRNA network, the hub lncRNAs were analyzed by using the cytoHubba ^27^ plugin in Cytoscape. We defined the hub lncRNA threshold as edges ≥ 3. The Cox proportional hazards regression model was employed to analyze the hub lncRNAs in the ceRNA network. The survival periods of the glioma patients were obtained from TCGA. Statistically significant lncRNAs affecting the survival period (P < 0.05) were determined by Cox regression univariate analysis to construct Kaplan–Meier survival curves for patients with glioma.

### 2.2 Prediction and evaluation of small open reading frames (smORFs) encoding TM peptides

The complete sequences of the lncRNAs were obtained from the NCBI Nucleotide database (https://www.ncbi.nlm.nih.gov). A combination of the above ceRNA network and four bioinformatic methods was used to predict the encoding ability of TM peptides in lncRNAs: 1) The candidate lncRNA sequences were predicted by Open Reading Frame Finder in NCBI (https://www.ncbi.nlm.nih.gov/orffinder/) by using the nucleotide sequences above to find smORFs. 2) The Coding Potential Calculator (CPC) and CPC2 ^28 29^ were used to access the quality of ORFs. 3) The BLAST tool in NCBI was used to detect the conservation of lncRNA sequences between different species. 4) The translated amino acid sequences of the smORFs were then used to identify the potential TM regions via TMHMM Server v2.0 ^30^ based on the hidden Markov model.

### 2.3 Peptide modeling and MD simulation

#### 1) Preparation of hub lncRNA-encoded candidate small peptide models

We prepared candidate small peptide models by using I-TASSER ^31^ online software. The I-TASSER server was developed for protein structure predictions with homology modeling based on the ab initio algorithm, and it was ranked as the best automated method for protein structure prediction in the Critical Assessment of Structure Prediction (CASP) experiments ^32 33^. Because all of the small peptides contained potential α-helix TM regions, we marked the TM regions and used the nuclear magnetic resonance (NMR) models of SLN and PLB as the reference structures for modeling.

#### 2) Preoptimization of peptide models

To select the most accurate small peptides for further simulation, the models predicted by I-TASSER to have the highest scores were chosen. The small peptides were embedded in a pre-equilibrium palmitoyl-oleoyl-phosphatidylcholine (POPC) membrane using the DESMOND software package ^34^. Then, the peptides were optimized by a 20 ns MD simulation. All of the parameters used were the same in subsequent simulations. Finally, the Rampage server was used for evaluating the models and the structure with best scores.

#### 3) Preparation of the peptide pentamer-membrane models and MD simulation

Two peptide structures were obtained from the prediction described above. To build the structure of self-assembled pentapeptides in the membrane, the PLB pentamer in membrane ^35^ (PDB code= 1ZLL) was used as the template due to its high similarity in the TM region. The equilibrated POPC membrane was used to mimic the SR membrane environment. The simulation was performed at a 14× 14×10 nm ^3^ box with three-dimensional periodic boundaries. The box was filled with a TIP3P ^36^ water model and used as water molecules. The size of the whole system was ∼140,000 atoms.

#### 4) MD simulation

To obtain the equilibrated initial pentameric peptides in the POPC membrane, the CHARMM36 force field was chosen for MD simulation. The particle mesh Ewald (PME) algorithm was applied for electrostatic interactions with a nonbonded interaction at 1.2 nm and a cutoff at 1.2 nm. The linear constraint solver (LINCS) algorithm was used to constrain the bond lengths. The system temperature was maintained at 310 K by a Nose–Hoover thermostat, corresponding to body temperature. The pressure was maintained semi-isotropically at 1 bar on the x and y axes by using a Parrinello–Rahman barostat, and the time step for simulation was 2 fs. A 20 ns equilibration process was applied under both constant volume (NVT) and constant pressure (NPT). Then, both peptides were simulated for 1 µs MD.

#### 5) Umbrella sampling and the potential of mean force (PMF)

The PMF of the two peptide systems was calculated by using the method described by Zhu et al ^37 38^. For all of the pentameric peptide systems, the H_2_O molecules were located in the cytoplasm bulk water region. The PMFs of water molecules were calculated from the two peptide models. One H_2_O molecule was pulled smoothly along the z-coordinate axis (the bilayer orientation was normalized to the z-axis), which corresponds to the direction from the N- to the C-terminal. An approximately 5 ns simulation was performed to obtain snapshots of the umbrella sampling windows. Seventy umbrella sampling windows for H_2_O molecules at an interval of 0.1 nm along the z-coordinate axis were produced to calculate the PMF from 3.5 to -3.5 nm (0 nm means the center of the membrane thickness). The H_2_O molecules were harmonically restrained by a force constant of 3000 kJ mol/nm^2^ on the z-axis. The simulation time for each window was 3 ns; the first 1 ns was used for system equilibration, and 2 ns was applied for analysis. The weighted histogram analysis method (WHAM) was used to compute the PMFs. The profile was generated by the GROMACS protocol ‘g_wham’ ^39^. Bootstrap analysis (N=50) was used to estimate the statistical error.

Both VMD and PyMOL software ^40 41^ were used to visualize all simulation processes. The GROMACS 5.05 software package ^42^ was used on the NVIDIA CUDA acceleration workstation and the Tianhe I Supercomputer at the National Supercomputer Center in Tianjin with Xeon processors. The total simulation times were ∼ 4 µs.

## 3. Results

### 3.1 ceRNA network and hub lncRNA selection

#### 1) Identification of DEmRNAs, DEmiRNAs and DElncRNAs

We analyzed lncRNA expression profiles in glioma patient tissues (n=700) and adjacent nontumor tissues (n=5). There were 420 downregulated and 128 upregulated lncRNAs within 548 aberrantly expressed lncRNAs (absolute FC>2, P<0.01; the volcano plot is shown in supplement Fig S1); and 2370 downregulated and 1615 upregulated mRNAs within 3985 aberrantly expressed mRNAs. Filtering analysis with the same criteria also identified 62 miRNAs differentially expressed between glioma and normal tissues, of which 20 were downregulated and 42 were upregulated (the full list of the RNAs can be found in the supplement Excel files).

#### 2) The glioma ceRNA network

Based on the ceRNA theory, the potential interactions among the above dysregulated genes have been predicted to understand the DElncRNA functions by bioinformatic analysis. Eight specific DEmiRNAs were identified to interact with 32 DElncRNAs through miRNA response elements. The miRTarBase and StarBase databases ^43 44^ were used to predict the candidate mRNA targets of the 6 DEmiRNAs and 41 mRNAs within the 3985 DEmRNAs that were used to build the ceRNA network. The ceRNA network involved 32 lncRNAs, 8 miRNAs and 41 mRNAs. Fig 1 shows the ceRNA network that was visualized by using Cytoscape software.

**Fig 1.**
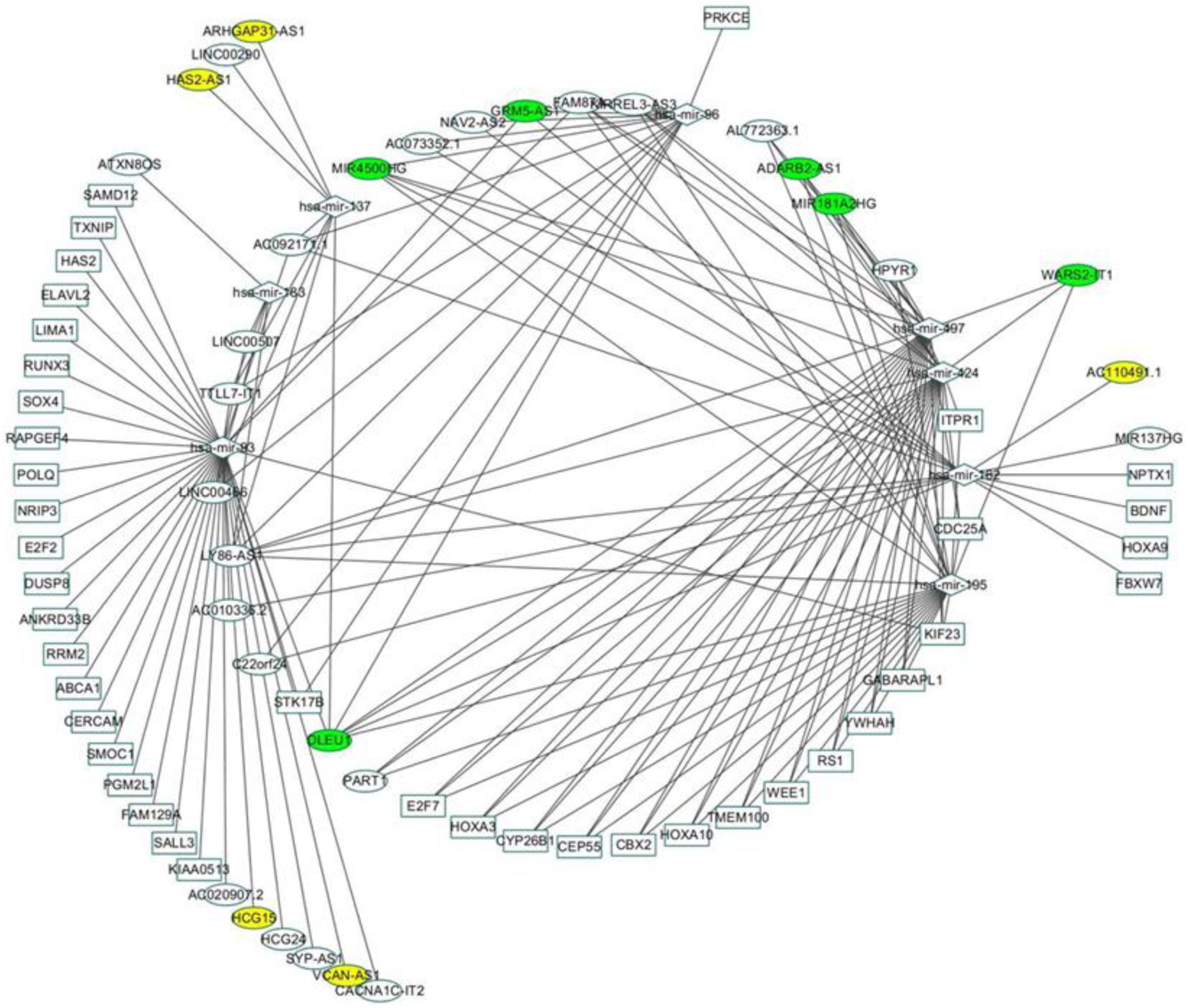
The ceRNA network. The green and yellow circles represent key lncRNAs and key hub lncRNAs (n >= 3), respectively.

#### 3) DElncRNAs and their associated clinical features

In analyzing the association between DElncRNAs and glioma patient survival periods, all DElncRNAs in the ceRNA network were chosen according to the bioinformatics analysis. The significant effects on survival were evaluated with *P* < 0.05 to identify the key hub lncRNAs with prognostic characteristics. In the ceRNA network, 11 lncRNAs (Table 1) were found to be associated with the overall survival of patients with glioma by univariate Cox regression analysis.

**TABLE 1.** The *P*-values of the lncRNAs associated with the survival rate of glioma patients.

| lncRNA | P-value |
| --- | --- |
| VCAN.AS1 | 9.99E-16 |
| ARHGAP31.AS1 | 3.00E-15 |
| AC110491.1 | 9.85E-14 |
| MIR4500HG | 2.04E-09 |
| HCG15 | 5.43E-05 |
| DLEU1 | 6.19E-05 |
| IT1 | 3.02E-04 |
| HAS2.AS1 | 6.77E-04 |
| GRM5.AS1 | 1.14E-03 |
| ADARB2.AS1 | 3.05E-02 |
| MIR181A2HG | 4.69E-02 |

The node lncRNAs were defined as edges of the ceRNA network >= 3. Finally, six node lncRNAs (MIR4500HG, GRM5-AS1 (Fig 1), ADARB2-AS1, MIR181A2HG, WARS2-1T1 and DLEU1) were found. Kaplan–Meier survival curves indicated that there are four node lncRNAs (MIR181A2HG, MIR4500HG, GRM5-AS1 and ADARB2-AS1) positively correlated with overall survival, but IT1 and DLEU1 were negatively associated with overall survival (the Kaplan–Meier survival curves of the 11 lncRNAs are shown in the supplement file).

#### 4) Functional analysis of lncRNAs negatively associated with overall survival

The functional analysis revealed a negative correlation with overall survival. There were two lncRNAs, DLEU1 and IT1, in the above ceRNA network that were enriched in 103 GO biological process categories. The GO biological processes of the above two lncRNAs were involved in ion channel activity (GO:0005216) and ion gated channel activity (GO:0022839). This finding suggests that the high expression of the two lncRNAs leads to a low survival rate in glioma patients, probably due to their abnormal biological processes involving ion or water permeability, which was also our main focus in this study.

### 3.2 ORF and TM region identification

Using the methods described in the Methods section, the candidate ORFs in hub lncRNAs were searched by ORF FINDER ^45^ online software [https://www.ncbi.nlm.nih.gov/orffinder]. We found that DLEU1 and IT1 contain ORFs. Then, the ORFs were translated into amino acid sequences and analyzed by TMHMM 2.0 to predict the TM regions. The CPC and CPC2 web tools were used to assess the quality of the lncRNA ORFs. Then, the BLAST tool in NCBI was used to compare the homology of DLEU1 and IT1 in order to determine whether they are conserved between different species. The following criteria were used: 1) The sequence must be a continuous ORF in a lncRNA. 2) The amino acid sequence, including the TM region, has to achieve a 100% probability (supplement Fig S3). 3) The potential TM region has to be conserved and compared to those in other species by multiple sequence alignment in the NCBI database. 4) Given the length of the identified small peptides (SLN, PLB, MLN, DWORF), the number of amino acids in the predicted peptide must be =< 45. 5) The ORFs quality must be >95% as assessed by both CPC and CPC2. 6) To ensure reliable results, we repeated the above steps using the known lncRNA-encoded small TM peptides MLN, DWORF, SLN and PLB. Through the above steps, we identified two possible TM peptides, encoded by ORF1 and ORF8, in the lncRNA DLEU1 according to the conditions above; these peptides were suitable for subsequent functional study. The results are shown in supplement figures S2-S4.

### 3.3 Model construction and MD simulation

Our results showed that the lncRNA DLEU1 was closely related to tumor proliferation and migration, while IT1 had no candidate peptides encoding ORFS. The subsequent molecular modeling and MD simulation focused on DLEU1’s 2 ORFs. The two ORFs obtained from DLEU1 contained the candidates of the small TM peptides, and we built the secondary structure of the ORFs using the I-TASSER online software package based on the ab-initio algorithm. Five candidate small peptides were predicted for each ORF. The peptides chosen for MD simulation had higher C-Scores, and the TM region appeared to have an α-helical configuration. The modeled peptide was embedded in a pre-equilibrated POPC membrane, and then a 50 ns MD simulation for optimization of the pentameric assemblies was performed until the RMSD became smooth, and then the optimized structure was extracted. Homology modeling for the pentameric candidates of the small peptides was performed as we described previously for the SLN and PLB pentamer structures. Two small peptide pentamers were embedded into the membrane, then 1 µs (1000 ns) of MD simulation was performed for optimization, and the stability of the pentameric assemblies during the optimization process was evaluated using the C-α’s RMSD and the Rampage online software. As shown in Fig 1a and b, after ∼100 ns and ∼30 ns, the RMSD curves for ORF1 and ORF8 were flattened, respectively, indicating that the two models were stabilized within 1 μs. The RMSD of both peptide pentameric assemblies showed that the RMSD of the TM region was ∼0.1 nm less than that of the backbones, indicating that the TMs have a higher stability. Comparing the two pentamers, ORF1 showed a more flexible backbone than ORF8, and the RMSD of the ORF8 pentamer was ∼0.1 nm less than that of the ORF1 pentamer.

### 3.4 Computation of H_2_O and ion permeability

To quantify the energy barrier for an individual H_2_O molecule through the DLEU1 candidate peptide channels, free energy profiles were used to identify the individual water molecular permeation from the cytoplasm (-z valued side) to the lumen (+z valued side) region of each pentamer. Fig 3 shows the PMF of H_2_O permeating through the two pores. The z-coordinate was defined as a function of the length of the pore. A H_2_O molecule was pulled into the pentamer channel, and the energy barrier (PMF) decreased sharply from 0 to -5 kJ/mol between -3 to -2.5 nm along the z-coordinate. Then, the curve ascended slowly from -2 to 0 nm. After a stable PMF barrier from 0 to 1.5 nm, there was another descent. These profiles are attributed to the energetic cost of moving H_2_O through the pore. The maximum free energy barrier for H_2_O crossing the ORF1 pentamer pore was ∼5 kJ/mol at Z_M_=0 to 1.5 nm. For the ORF8 pentamer, when a H_2_O molecule permeated through the pore from -3.5 to 0 nm, the energy cost rapidly increased from 0 to 20 kJ/mol, which corresponded to the maximum energy barrier where three leucine residues were placed. The location of the leucine residue showed the strongest hydrophobicity of the pore. Subsequently, the free energy profile drops rapidly from 0 to 3 nm and eventually to zero. The simulation results indicated that the H_2_O molecule penetrated more easily through the pentameter pore of DLEU1 ORF1 than that of ORF8. The lower energy barrier for water transportation from the cytoplasm to the lumen suggested that the H_2_O molecules had a favorable flow in the ORF1 pore. The amino acid residues of ORF1 and ORF8 at Z_M_=0 and -1 are serine and leucine, respectively. This indicates that the side chains of serine and leucine have a greater influence on water permeability.

**Fig 2.**
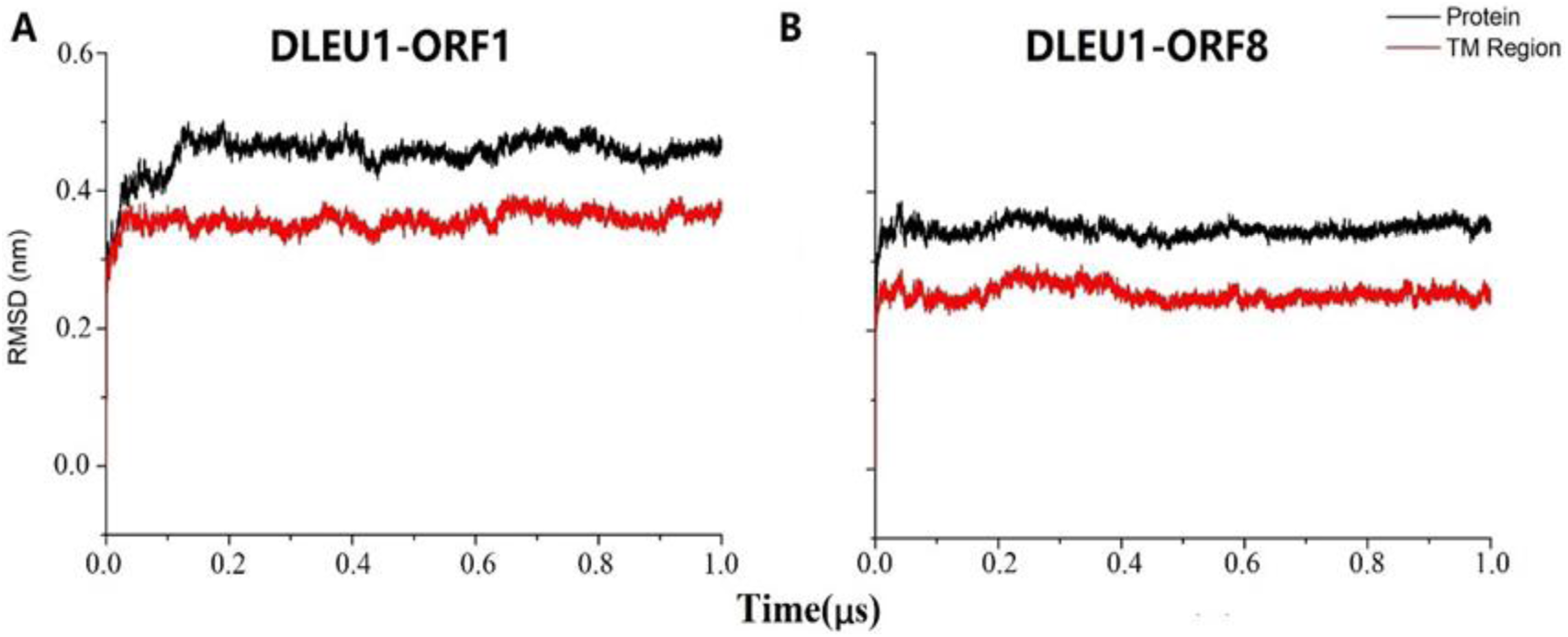
Structural stability of pentameric DLEU1 ORF1 (A) and ORF8 (B) peptides. The black and red solid curves represent the root mean square deviation (RMSD) of the whole protein and TM region, respectively.

**Fig 3.**
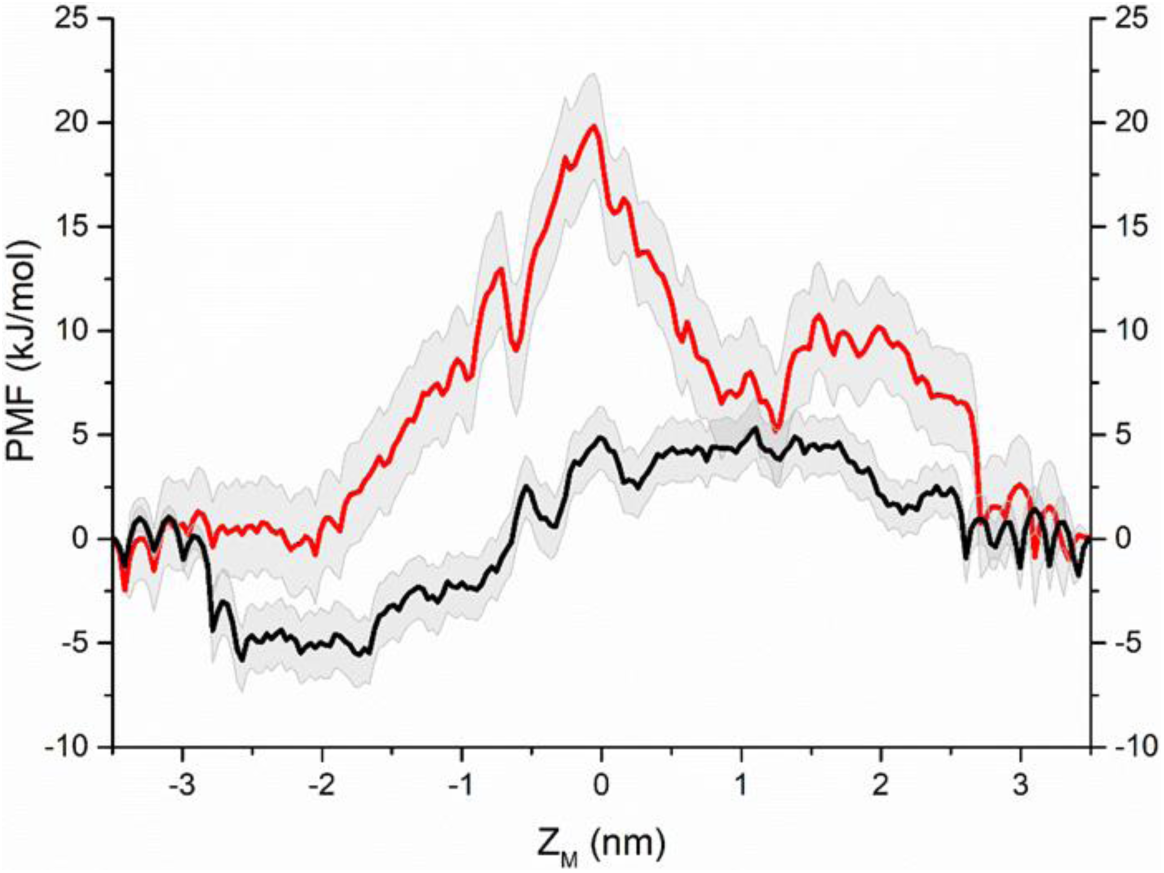
The potential of mean force (PMF) associated with water permeation through the channel pore of ORF1 (black curve) and ORF8 (red curve) pentamer.

Due to the substantial energy barrier difference in water permeation between the pentamer pores of ORF1 and ORF8, if there is an energetic barrier to water permeation, then it is likely that a gate will also be impermeable to ions. Therefore, we are interested in focusing on ORF1 pore’s ability to permeate ions. To check whether the ORF1 pentamer plays a role as an ion channel, the PMFs of Na^+^ and Cl^-^ through the ORF1 pore were calculated as well. Both Na+ and Cl-had to overcome a large energy barrier in the pore. As shown in Fig 4, for Na+, the maximum energy barrier reached 20 kJ/mol at Z_M_=0. The maximum energy barrier of Cl-was 25 kJ/mol at Z_M_=-1 nm, which was slightly larger than that of Na+. The energy barriers of Na+ and Cl-revealed that the ions were not allowed to freely permeate through the ORF1 pentameric channel.

**Fig 4.**
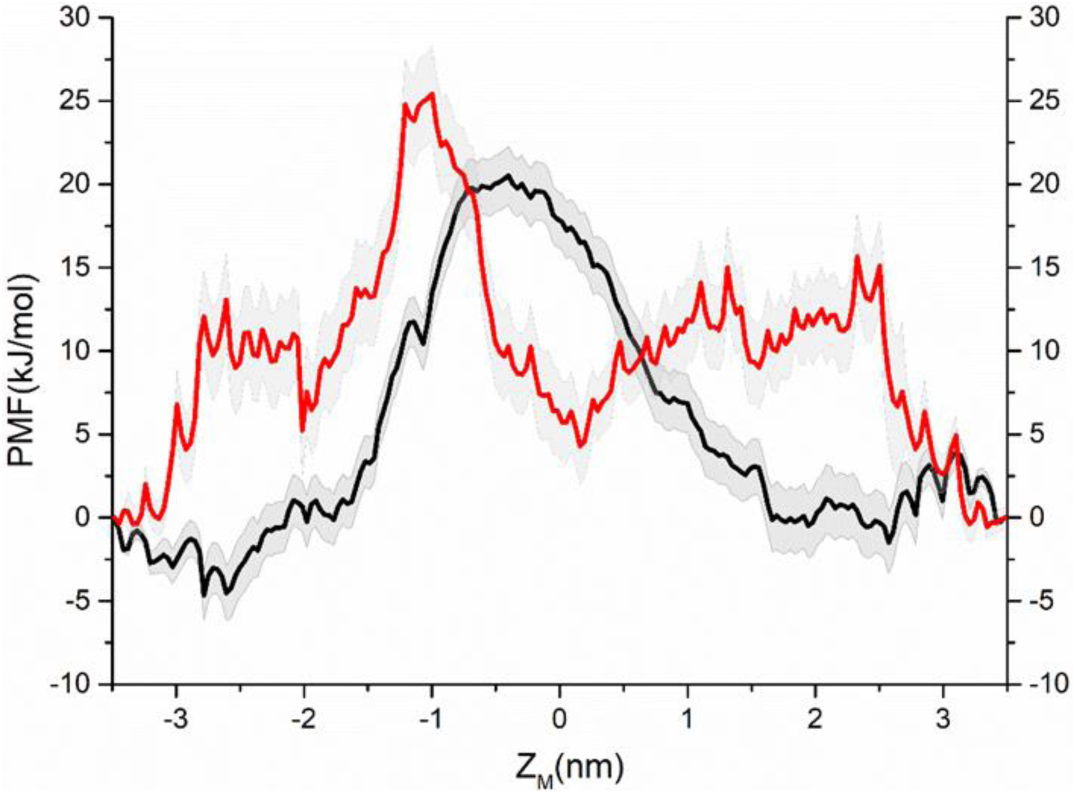
PMF associated with Na+ (black curve) and Cl- (red curve) permeation through the channel pore of the ORF1 pentamer.

The number of water molecules in the ORF1 pore showed complete water permeation and periodic variation during the 1 µs MD simulation (see Fig 5, red and gray curve). The quantity of water molecules changes between ∼7 to ∼20 kJ/mol at Z_M_= -1∼1 nm, and the time interval is ∼0.1 µs. The permeated number of Cl^-^ ions also fluctuated periodically as water molecules changed at the 0∼2 scale, indicating that the ORF1 pore presented a higher-energy barrier for Cl^-^ permeation. This result was consistent with the PMF profiles. Both the PMF and water molecular changes indicated that the ORF1 channel mainly functioned as a water channel rather than as an ion channel.

**Fig 5.**
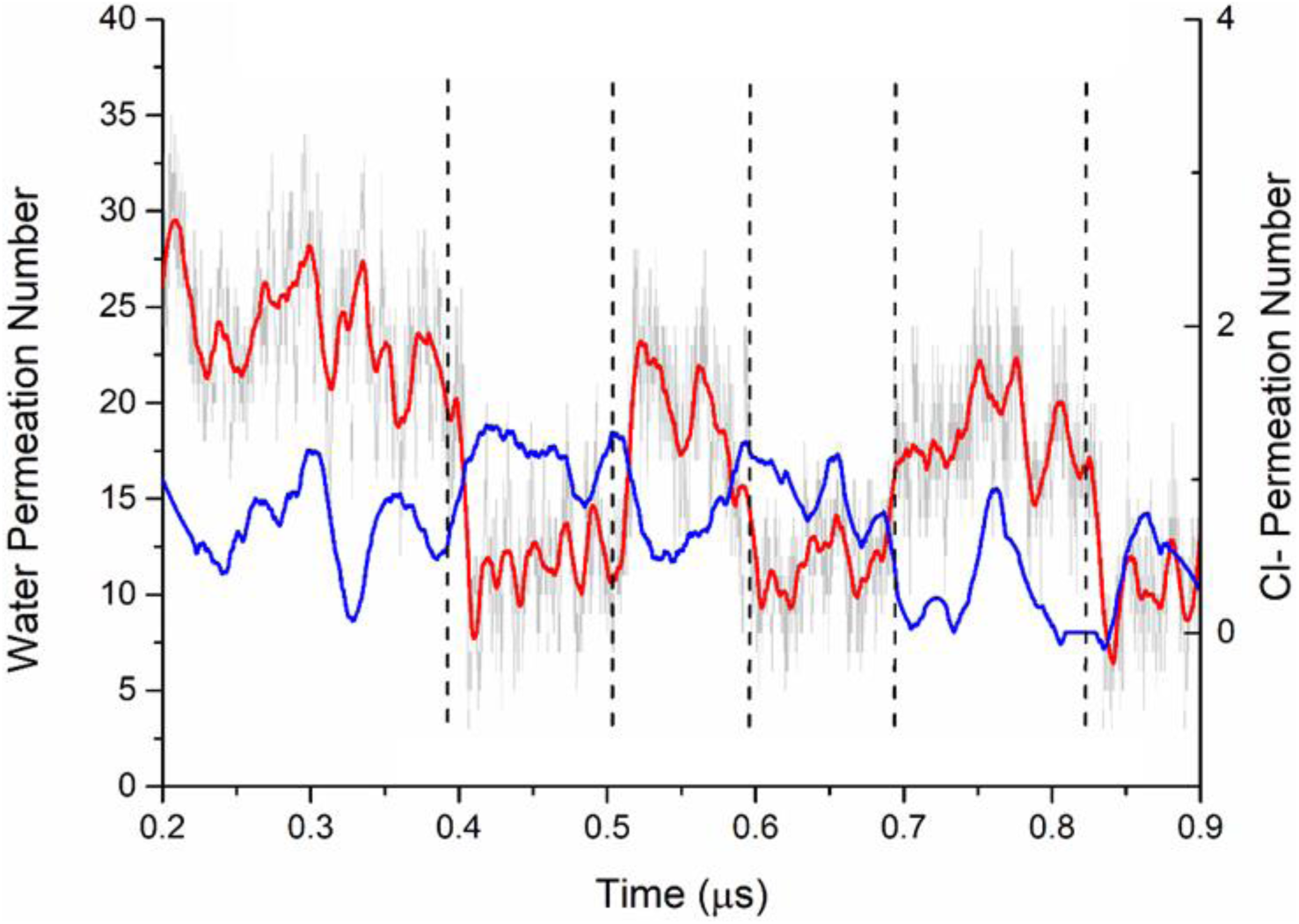
The change in ORF1 with water and chloride ions during the 1 µs simulation. The gray curve represents the absolute quantity of water molecules. The red and blue curves represent the smooth line of water and chloride ions, respectively (200 pts).

## 4. Discussion

Glioma is the most common intracranial tumor and is characterized by various abnormal gene-protein regulatory events that drive tumorigenesis ^46^. At present, many efforts have been made to elucidate the role of lncRNAs and the related membrane proteins involved in tumorigenesis, progression and cerebral edema ^47 48^. According to a previous study, lncRNAs can participate in the regulation of cellular ions and water by encoding small TM peptides, but the mechanism of small TM peptides is still unknown.

Currently, there are no accurate methods for predicting lncRNA-encoded small peptides. In this study, ceRNA network, survival, and GO enrichment analyses were conducted to identify hub lncRNAs with channel functions. Subsequently, various bioinformatic methods and tools were used to predict and validate the small TM peptides encoded in ORFs of lncRNAs in glioma. Finally, the ORFs were translated into amino acid sequences. A combination of ab initio modeling, molecular dynamics simulations and other methods were used to determine the pore characteristics of the pentamers.

To establish the ceRNA network among mRNAs, lncRNAs and miRNAs in gliomas, 705 sequencing data points in the TCGA database from a cohort of tissue samples (700 cancer tissues and 5 adjacent tissues) were characterized to establish a ceRNA network. We identified a total of 4,595 aberrantly expressed lncRNAs, mRNAs and miRNAs and successfully constructed a prognosis- and lncRNA-related ceRNA network in gliomas by biological prediction. Eleven lncRNAs with significant prognostic differences were detected (Table 1). Two of the lncRNAs, DLEU1 and IT1, were negatively correlated with the potential prognosis of glioma patients. The high expression of DLEU1 and IT1 significantly reduces the survival rate of glioma patients. In subsequent GO enrichment analysis, we found that the functions of the two lncRNAs were related to channel activity, especially in particle transport and TM transport ^49^. This finding indicated that their effect on prognosis might be related to the dynamics of intracellular ions and water, but there is no evidence that the lncRNAs are directly involved in cellular ion transport. Therefore, the ORFs of the lncRNAs were identified by a bioinformatic method, it could be translated into small TM peptides.

Many studies have shown that the permeability and movement of water and ions play a key role in regulating the migration/invasion of cancer cells ^50 51^, which has an important relationship with the cerebral edema caused by glioma. The movement of water may accelerate tumor migration, making tumors more prone to metastasis ^19^. Although water infiltration is often regulated by the classical AQP water channel family ^52^, lncRNA-encoded small TM peptide oligomers such as PLB and SLN may also play an important role. The water permeability of the small TM peptide oligomer channel works in the same manner as PLB pentamers in muscle cells. To further confirm the possible permeability of these peptide oligomer functions in gliomas, we chose DLEU1 for subsequent simulations. DLEU1 activates KPNA3, which is related to the increased proliferation and migration of cancer cells ^53^. Some reports have indicated that DLEU1 can increase cell proliferation; our results showed that ORF1 and ORF8 in DLEU1 could encode small TM peptides. Through molecular modeling, MD simulations, PMF computation and structural analysis, we found that there were different water and ion behaviors between the ORF1 and ORF8 pentamers.

The 1 µs MD simulation results of the two peptide pentamers (ORF1 and ORF8) showed that the RMSD was stabilized within 0.1 µs. The RMSD of the whole protein and TM region fluctuations were maintained within 0.5 nm. Both peptide RMSD and simulation trajectories showed that there were no structural fractures or depolymerizations, indicating that the peptide pentamer channels remained stable in the simulation environment. This result was consistent with the simulation results of our previous SLN and PLB pentamers ^54^. In comparing the ORF1 and ORF8 pentamers, the RMSD of the ORF1-encoded peptide pentameter was ∼0.1 nm higher than that of the ORF8-encoded peptide pentamer, suggesting that the ORF1 pentameter was more flexible than the ORF8 pentamer.

The water permeability of the ORF1 pentamer channel (black curve in Fig 3) can be characterized as a three-stage process: i) the spontaneous water molecule entry stage with -5 kJ/mol free energy; ii) the overcoming stage of a ∼8 kJ/mol energy barrier at the central region of the channel pore at approximately Z_M_=-2 - 0 nm; and iii) the overcoming stage of the decreasing energy barrier from ∼3 kJ/mol. The energy barrier of the ORF8 channel was completely different from that of the ORF1 channel. When water molecules entered the ORF8 pores, the energy barrier rose steeply until the maximum energy barrier was ∼20 kJ/mol, which was nearly 15 kJ/mol higher than that of ORF1. Then, the curve dropped steeply, experiencing a flattened barrier region until the water molecule permeated through the pore. Water molecules can spontaneously permeate through the ORF1 pentamer channel pores, but not through the ORF8 pentamer channel due to the higher-energy barrier. The count diagram (Fig 5) of water molecules also showed that the water permeation through the ORF1 channel was a spontaneous process. The quantity variation in water molecules showed an obvious hydrated-dehydrated alteration, suggesting that the pentameric ORF1 channel was a hydrophobic channel. However, the counting diagram of Cl^-^ showed no spontaneous TM activity through the pore. Interestingly, the distribution of water and Cl^-^ along the channel pore showed competing relationships with each other. Cl^-^ ion occupation in the pore could block water permeation. To further validate the ionic function in the ORF1 pentamer, we calculated the PMF of the Na^+^ and Cl^-^ permeation through the ORF1 pentameric channel. The PMF results suggest that the highest energy barriers for Na^+^ and Cl^-^ were 20 kJ/mol and 25 kJ/mol, respectively. The MD trajectories showed that neither Na^+^ nor Cl^-^ pass through the pore. Intriguingly, Na^+^ ions exhibited Brownian motion in water without entering the pores, but Cl^-^ ions appeared at the entrance of the pore, appearing to be ‘stuck’ at leucine amino acids and blocked by the channel. This finding was consistent with the results of the PMF and water distribution. We could see that the ORF1 pentamer channel showed water transport capacity but no Cl^-^ transport process. According to the permeation of water and ion simulation results, we could reduce the energy barrier for water and ion transport by placing the charge group at the mouth of the channel pore, such as that in the ORF1 peptide. This finding was also consistent with previous studies on water transport through narrow or nanopores ^55 56 57^. In addition, the hydrogen bond distribution between the water molecules and amino acids inside the pores of the ORF1 and ORF8 pentamers were different. The water molecules in the ORF1 pore have fewer hydrogen bonds than those in the ORF8 pore, which could lead to the difference in water permeation.

To reveal the difference in the water and ionic permeability of the small peptide pentamer channels of the two ORFs, the radii of the two pores were calculated as shown in Fig 6. These two ORF channels appeared to have very similar pore radius profiles. A minimum radius of ∼0.3 nm was larger than the van der Waals radius of the water molecule (0.19 nm), as shown in Fig 6C. However, the energy barriers of water permeation were quite different between the ORF1 and ORF8 pentamers (Fig 3). The structure and sequence of the two ORFs’ small peptides were analyzed. The results showed that the water permeability difference between the two ORFs’ pentamers was mainly affected by the following factors. (1) Although both the ORF1 and ORF8 small peptides have a valine\leucine\isoleucine zipper domain, ORF1 missed one residue of valine\leucine\isoleucine at the TM region (Fig 7 C). This weakened the intermolecular interaction among the small peptides of the ORF1 pentamer in the formation of the pentameric assembly. Consequently, these weakened interactions led to a more flexible pentameric structure and the easier permeation of water molecules. (2) There were four hydrophilic amino acids for ORF1 located in the N-terminal of the peptide (cytoplasmic region), while for ORF8, they were located in the C-terminal (lumen region). This caused a significant absolute energy difference between the water permeation of the ORF1 and ORF8 pores. (3) There were two positively charged groups (Arg2 and Arg3) at the N-terminus of the ORF1 peptides (Fig 7). These electric charges in arginine attracted water molecules, supporting the explanation of water accessibility on the N-terminus of the pentamer. Some water molecules spontaneously entered into the pentamer channel pore, while the Cl^-^ ions were blocked from the pore. In contrast, the N-terminus of ORF8 had no charged amino acid residues. Water molecules are hard to aggregate, reducing the probability of spontaneous permeability.

**Fig 6.**
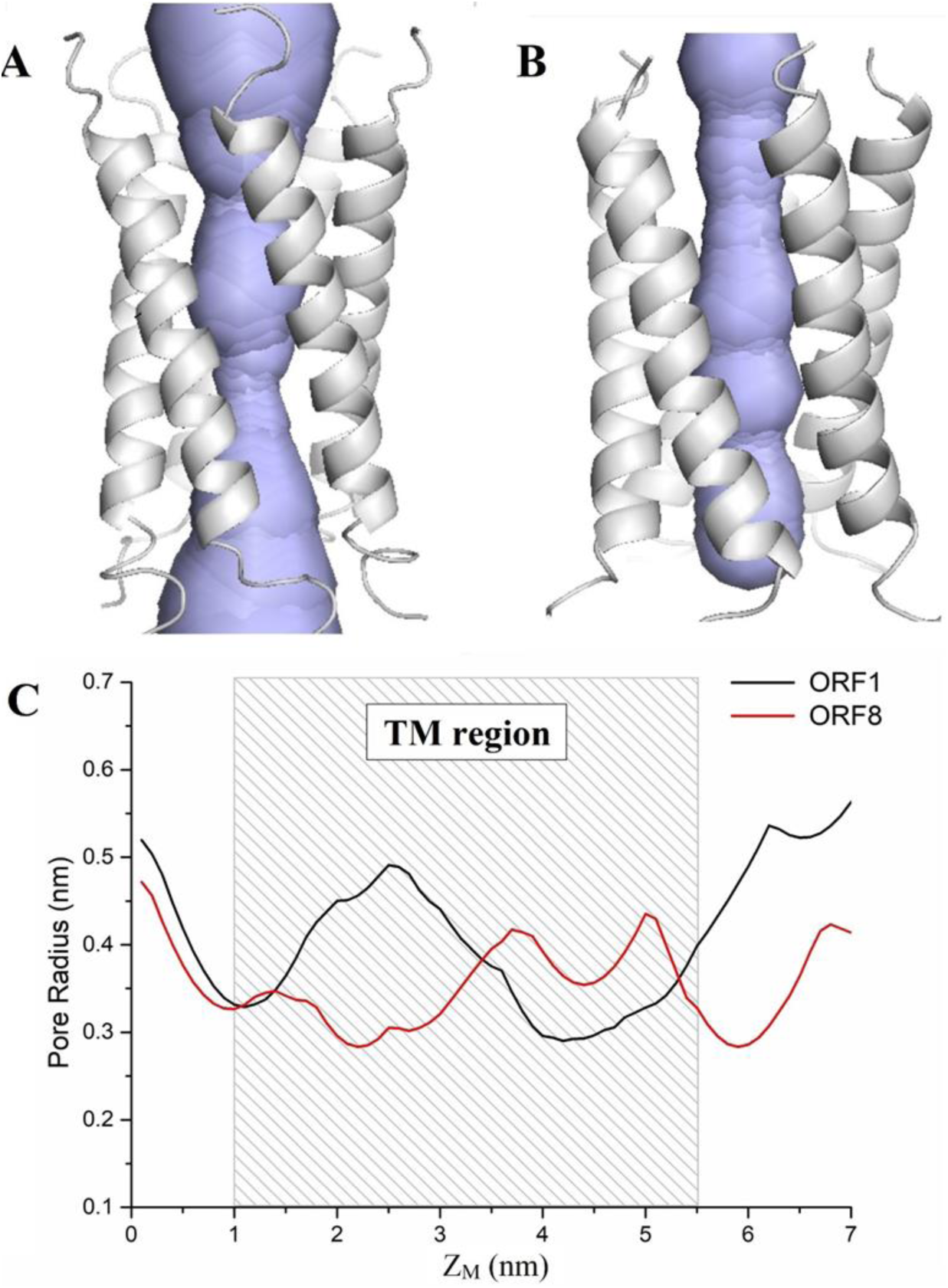
The radius of the ORF1 and ORF8 channel pores. A and B represent ORF1 and ORF8, respectively, with the pore’s lining surfaces in lavender. The rectangle with a sparse pattern in C is the TM region of the pore. The radii of the ORF1 (black) and ORF8 (red) pores are shown in C.

**Fig 7.**
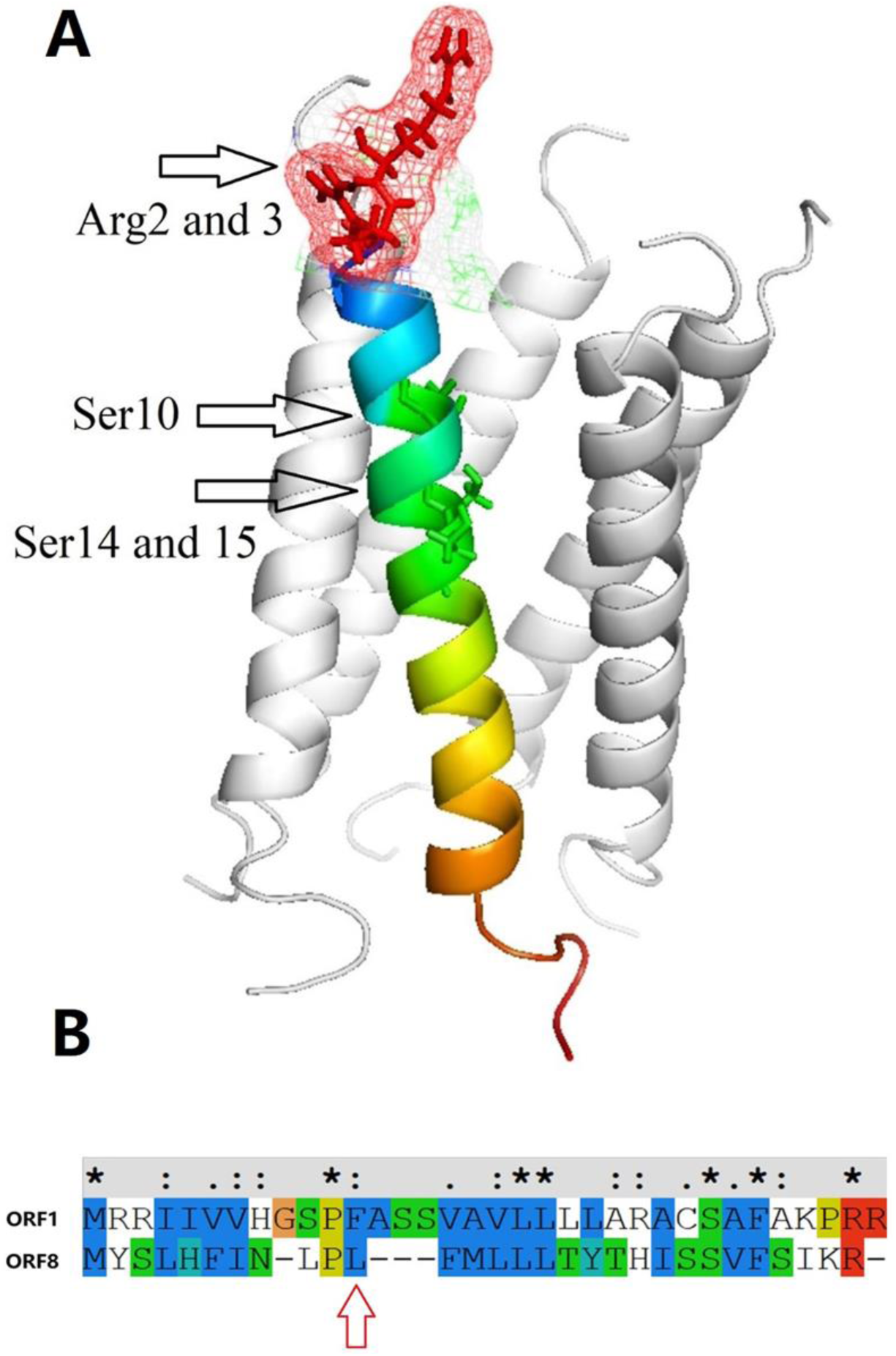
A, The pentameric ORF1 channel. The arrows indicate two charged amino acid groups, Arg2 and 3, and three hydrophilic groups, Ser10, 14 and 15. B. The sequence alignment of ORF1 and ORF8. There is one leucine in ORF8, but not in ORF1.

As some ORFs in lncRNAs can encode small peptides with biological functions ^17^, our results suggested that the ORFs of the hub lncRNAs with survival differences, including ORFs encoding small TM peptides, might affect the prognosis of glioma patients. The ORF1-encoded peptides can interact with each other on the membrane to form stable oligomeric hydrophobic channel pores. The ORF channel enhanced water influx, leading to water and ion leakage. The leaked ions and water would affect the microenvironment of the cell or interact with the known water and ion channel permeation. The intracellular and extracellular water and ion exchange causes tumor cell proliferation and migration, aggravating the occurrence of cerebral edema. Many small TM peptides have a general hydrophobicity that allows water and ions to influx. In this study, the simulations of the ORF1 and ORF8 pentamer channels showed significant differences in water and ion permeability, indicating that the hydrophobicity of the lncRNA-encoded small peptides was different. In addition, this difference explained why lncRNAs had different effects on cancer prognosis even though they had similar ORFs that could encode small TM peptides. The survival differences caused by lncRNA-encoded small TM peptides are very worthy of further study and corresponding experimental confirmation.

## 5. Conclusion

A model was built from a glioma ceRNA network to screen for differentially expressed hub lncRNAs and to identify ORF fragments encoding small TM peptides by bioinformatics methods and ab-initio homology modeling combined with MD simulation. Our established models revealed that the small peptides of the two encodable ORFs (ORF1 and ORF8) of the DLEU1 lncRNA could form pentameric channels. The simulations of the water behavior along the pentameric channel showed that the ORF1-encoded channel was possibly a hydrophobic gate. These findings were consistent with our two previous works of lncRNAs encoding the small peptide pentameric structures of SLN and PLB. We proposed that DLEU1 has the ability to encode small peptides with ion channel activity that might lead to an increase in glioma permeability, which in turn increases brain edema and even increases the risk of cancer cell development, invasion and metastasis. This mechanism could be used for prospective studies of the function of lncRNA encoding small TM peptides and for the development of new glioma treatment strategies.

## Supporting information

all supplemental table

## Acknowledgments

This study was supported by the National Natural Science Foundation of China (NSFC-31900894) and The Science & Technology Development Fund of Tianjin Education Commission for Higher Education (2018KJ067).

## Notes

### Competing Interest Statement

The authors have declared no competing interest.

