## Supplementary material for "Prediction of LncRNA Encoded Small Peptides in Glioma and The Oligomer Channel Functional Analysis Using in Silico Approaches": all supplemental table: supplement figure.docx

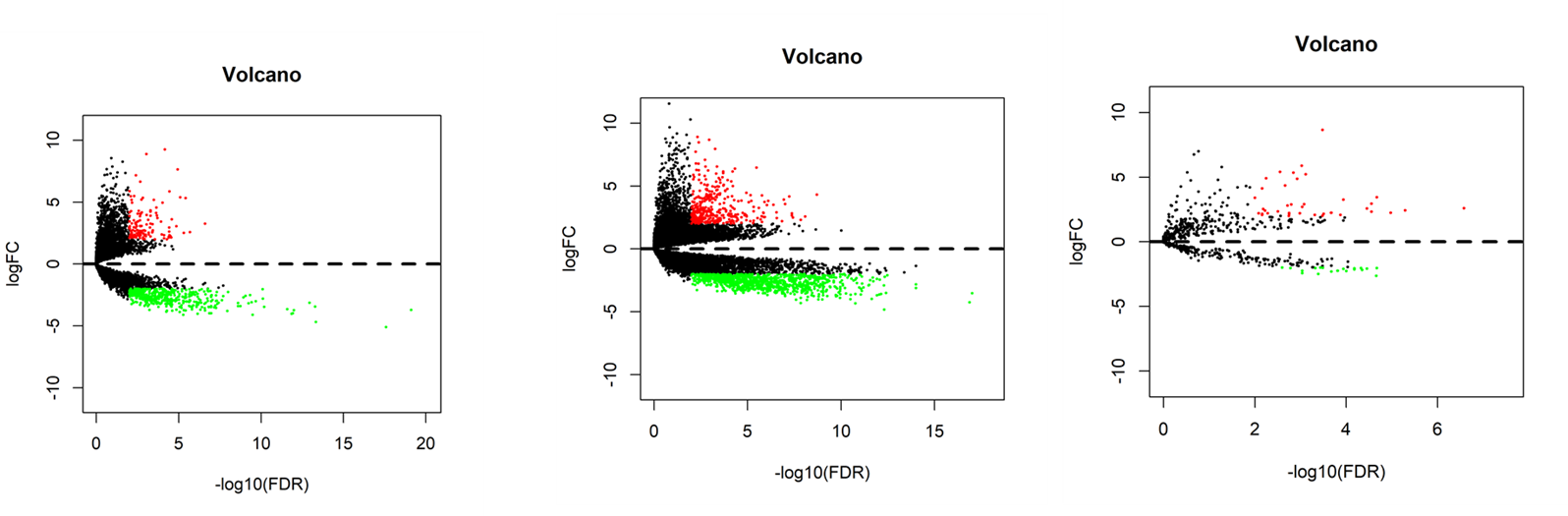

**lncRNA	mRNA	miRNA**

Fig S1. The volcano of lncRNA, mRNA and miRNA. The red and green dots represent upregulate and downregulated RNAs respectively.

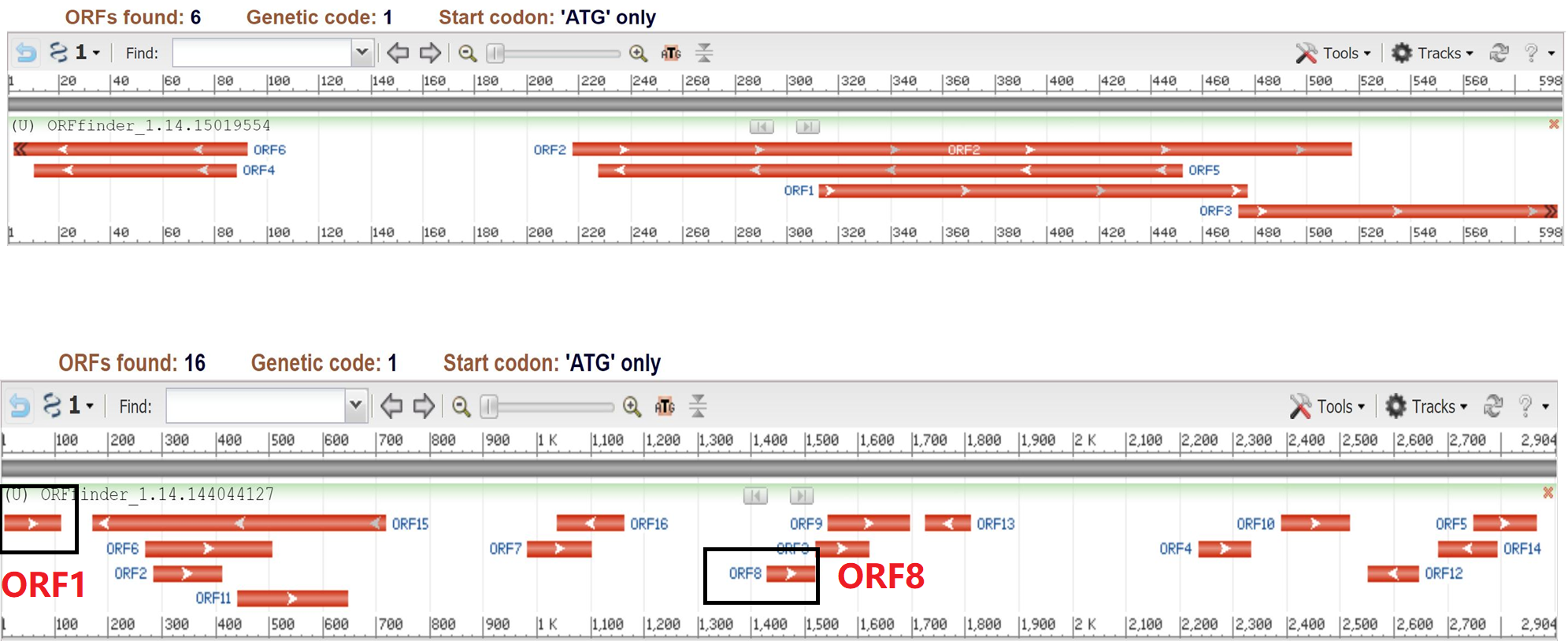

Fig S2 The predicted ORFs in WARS2.IT1 and DLEU1 lncRNA . the DLEU1 contain ORF1 and ORF8 can encoding small peptides.
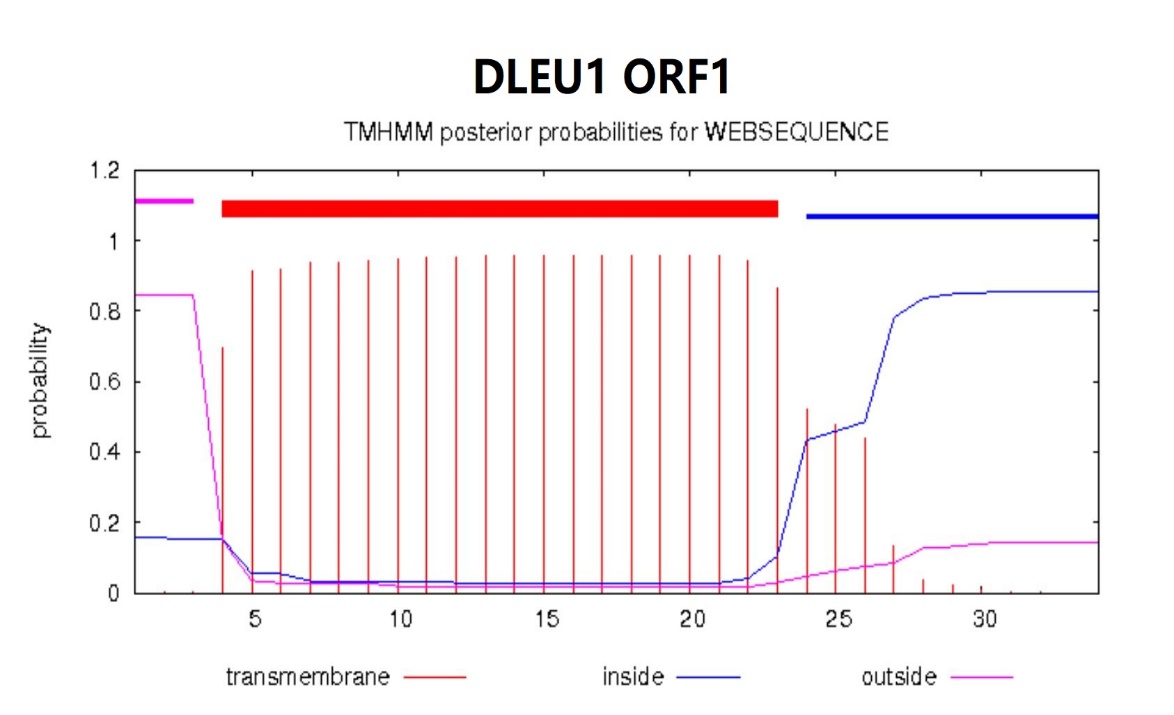

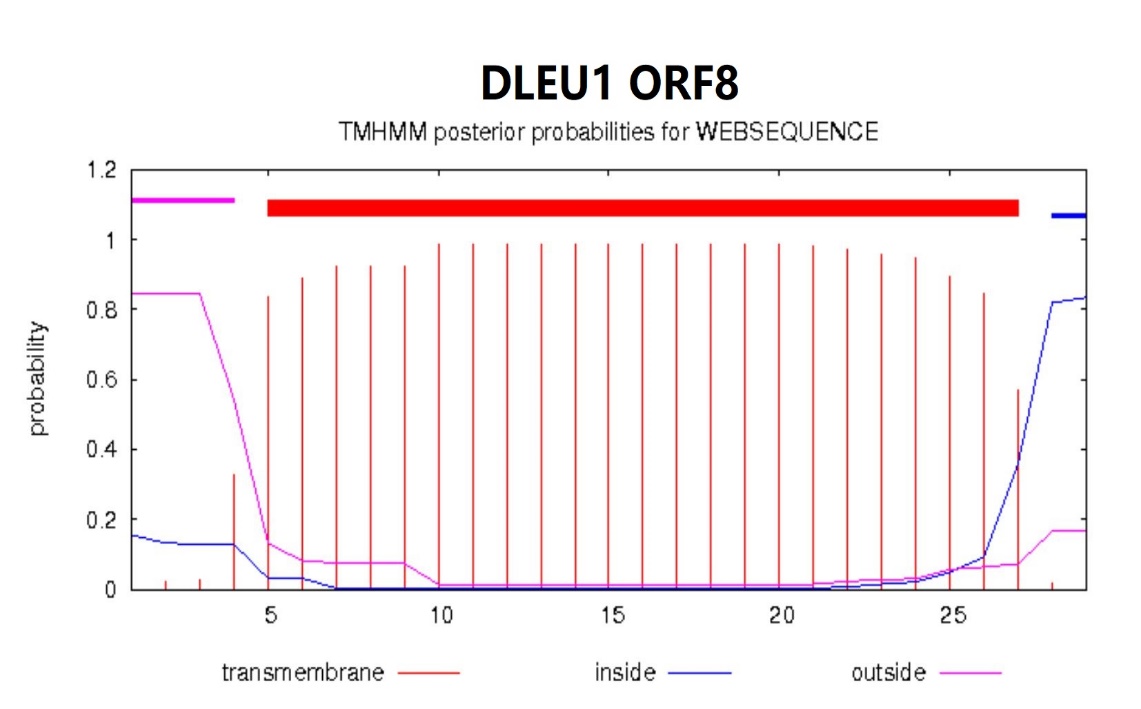

Fig S3, Two ORFs second structure, the red lines represent a predicted transmembrane α-helix configuration.

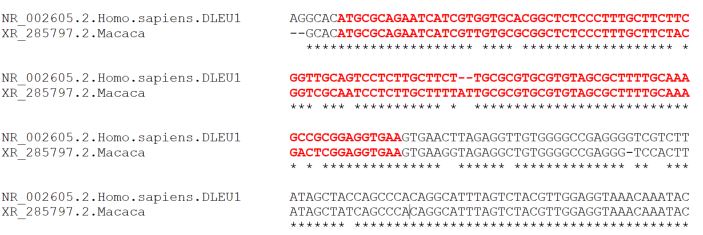

DLEU1 ORF1

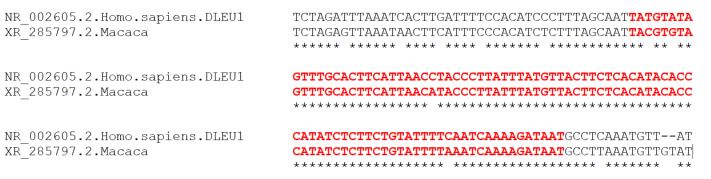

DLEU1 ORF8

Fig S4, Comparison of Two ORFs sequence conservation of DLEU1.

DLEU1 ORF8
