## Supplementary figures and images for "Prediction of LncRNA Encoded Small Peptides in Glioma and The Oligomer Channel Functional Analysis Using in Silico Approaches"

### Kaplan–Meier survival curves.docx

The 11 hub lncRNA’s Kaplan–Meier survival curves

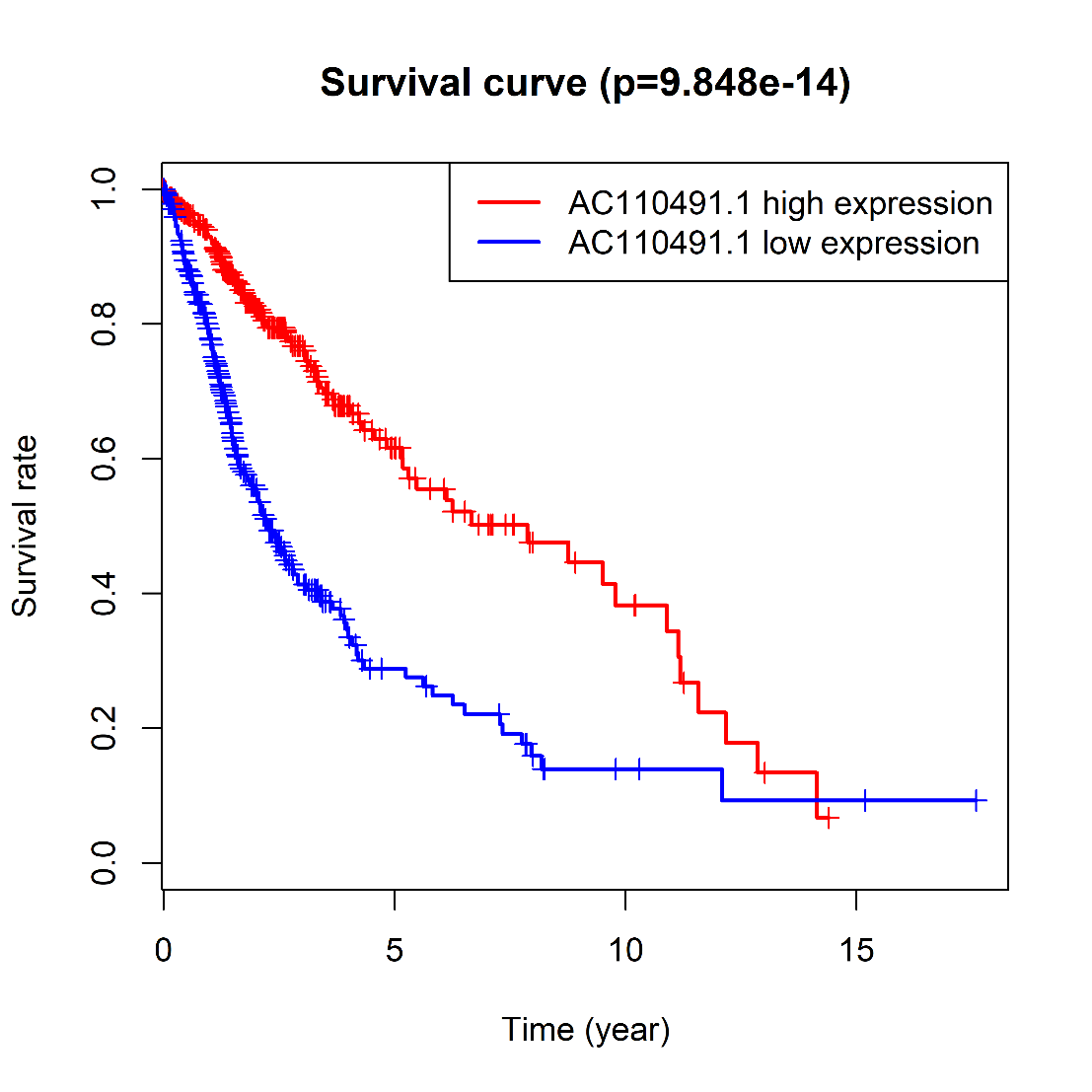

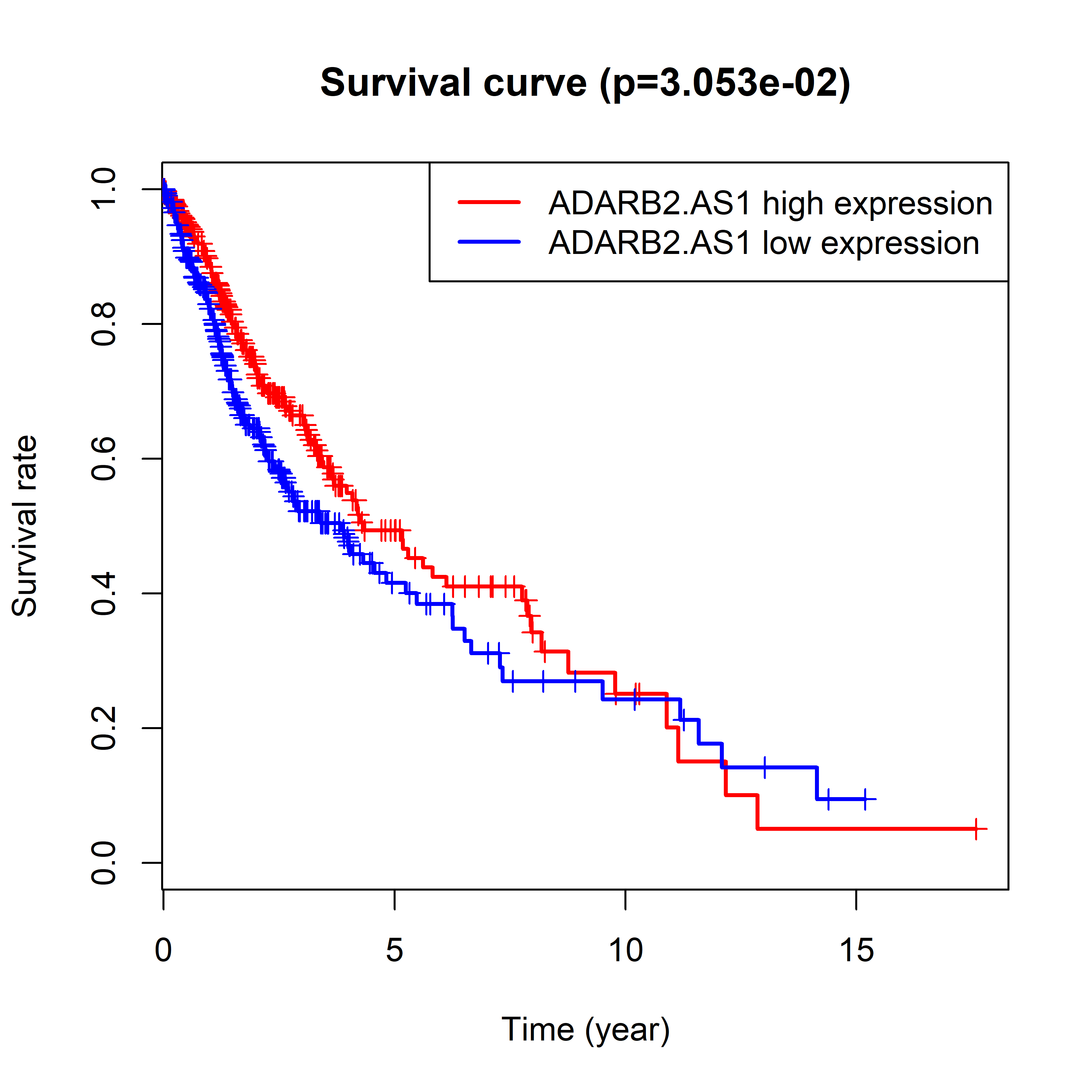

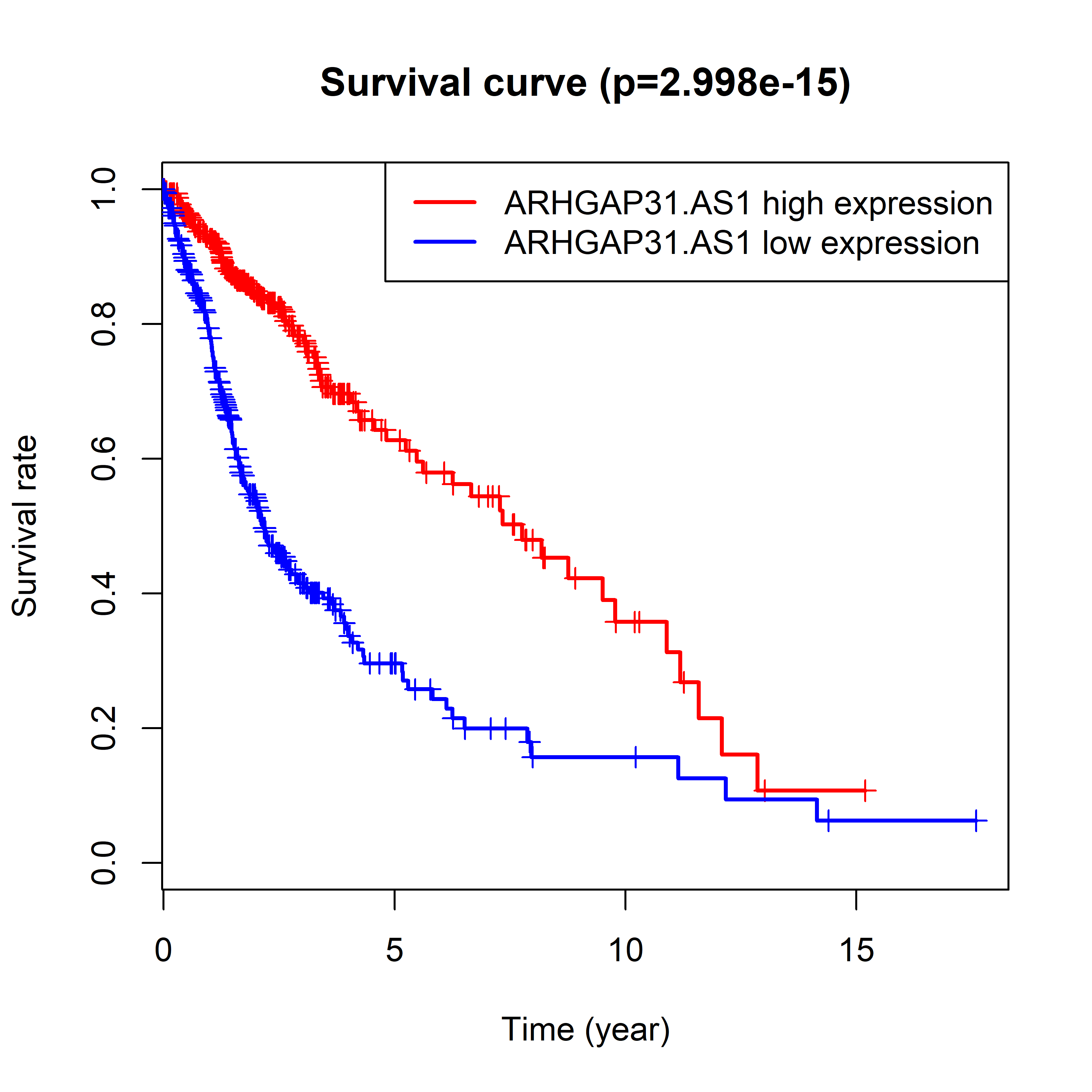

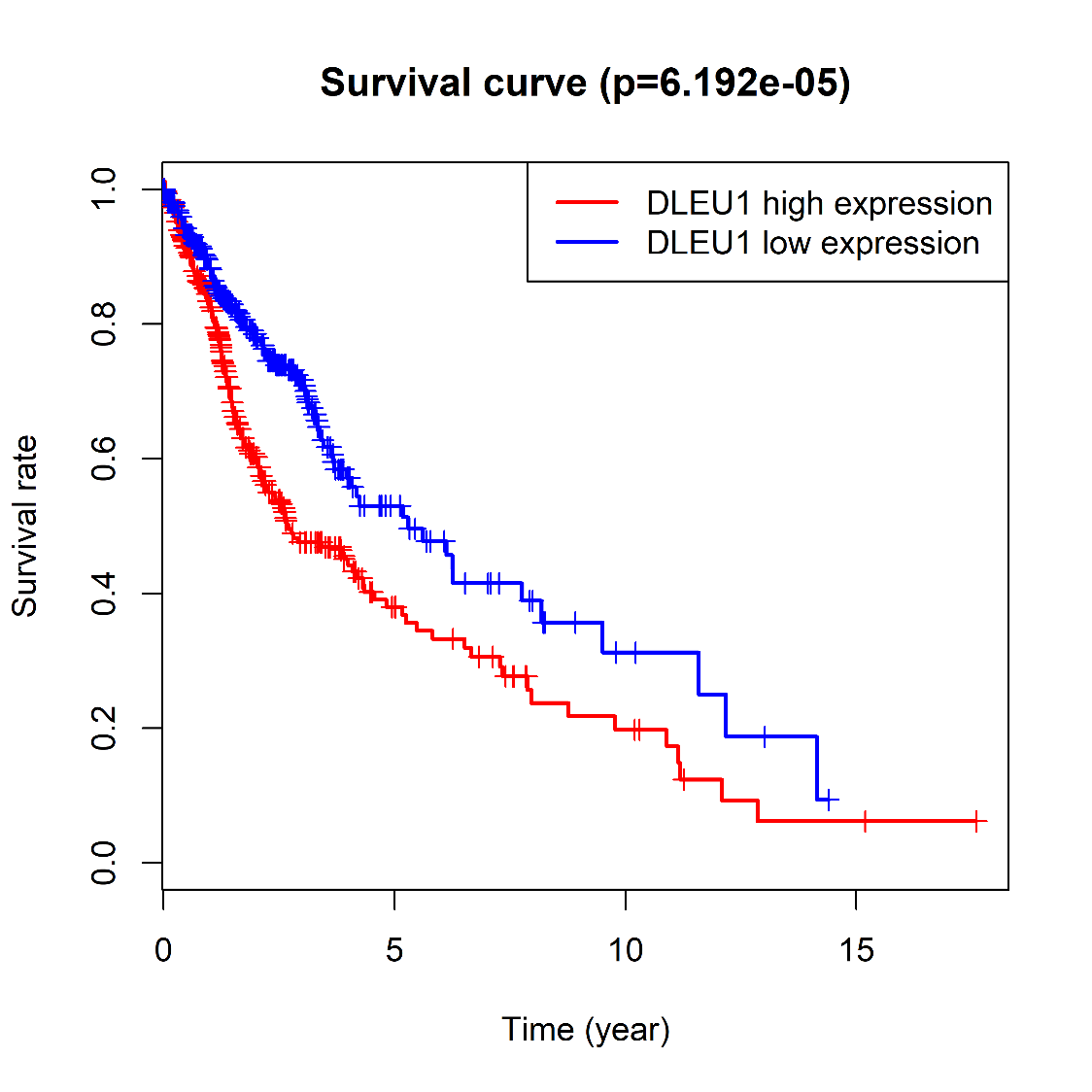

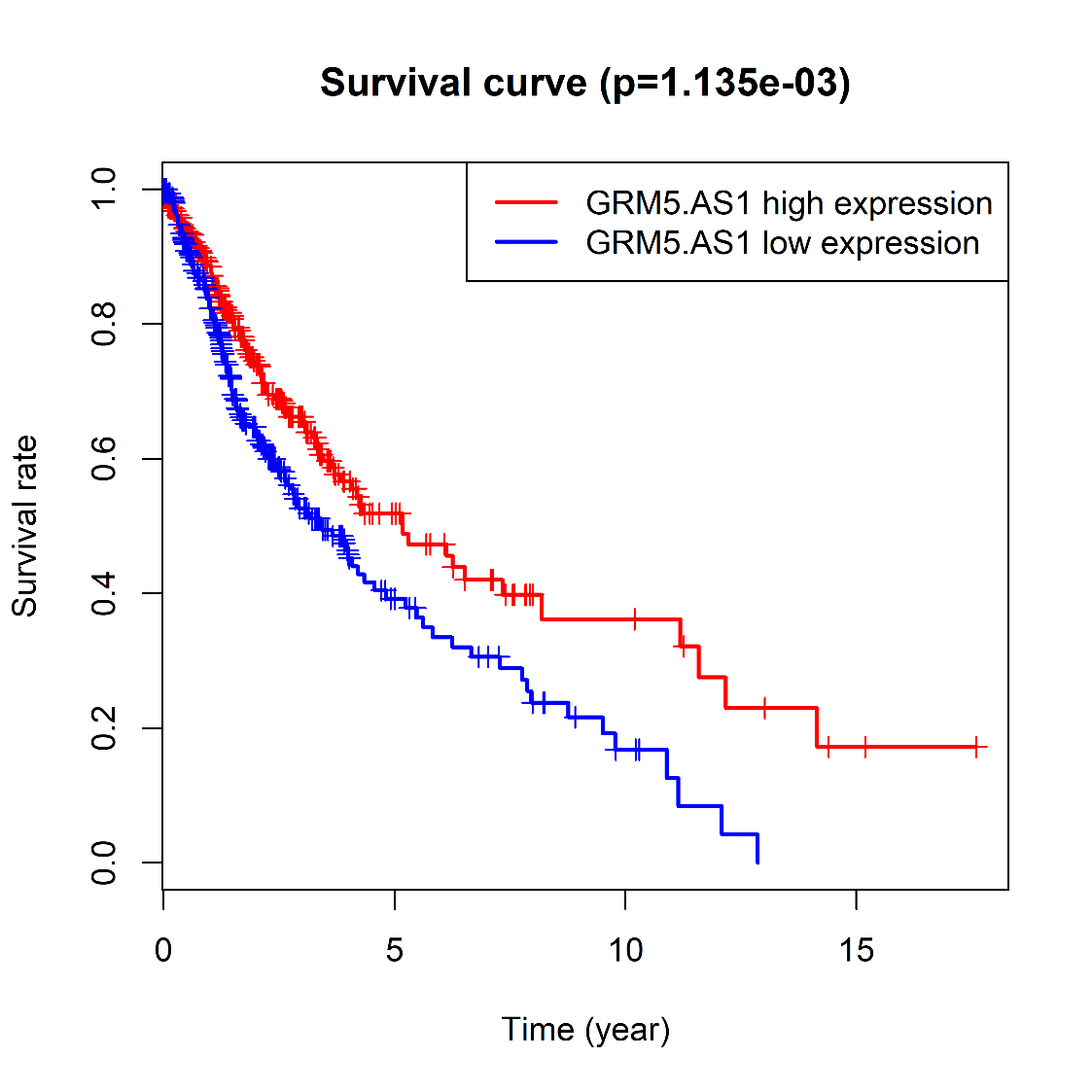

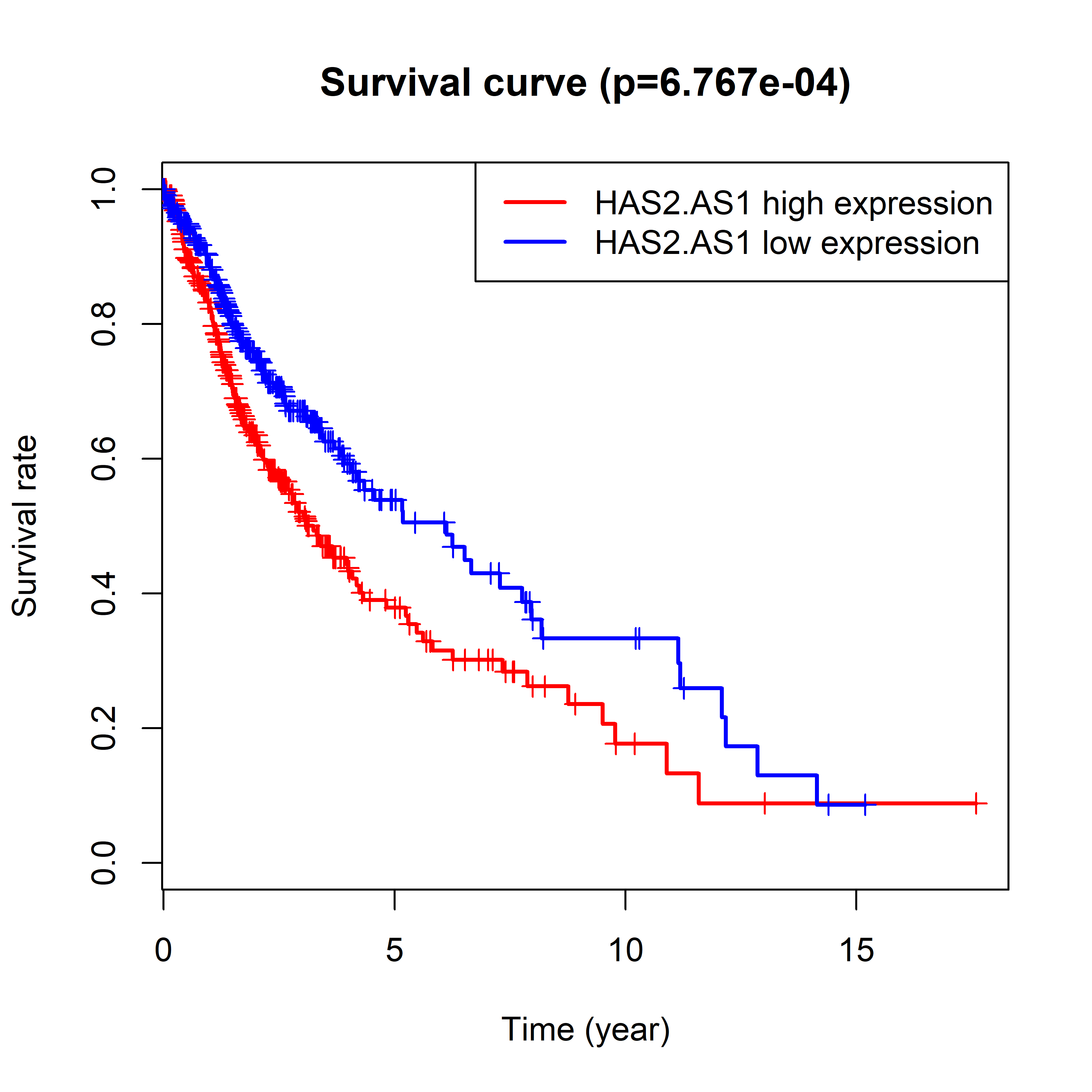

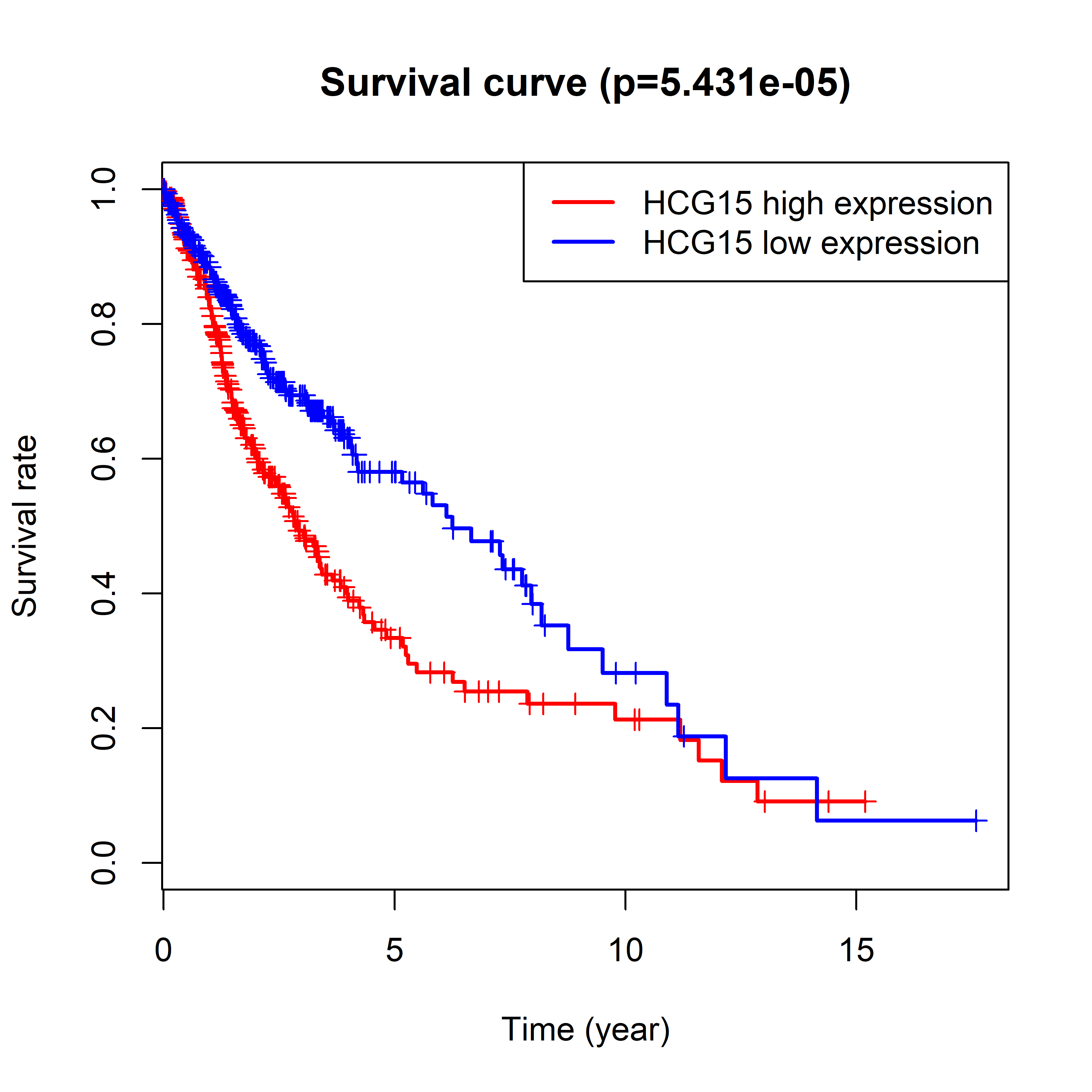

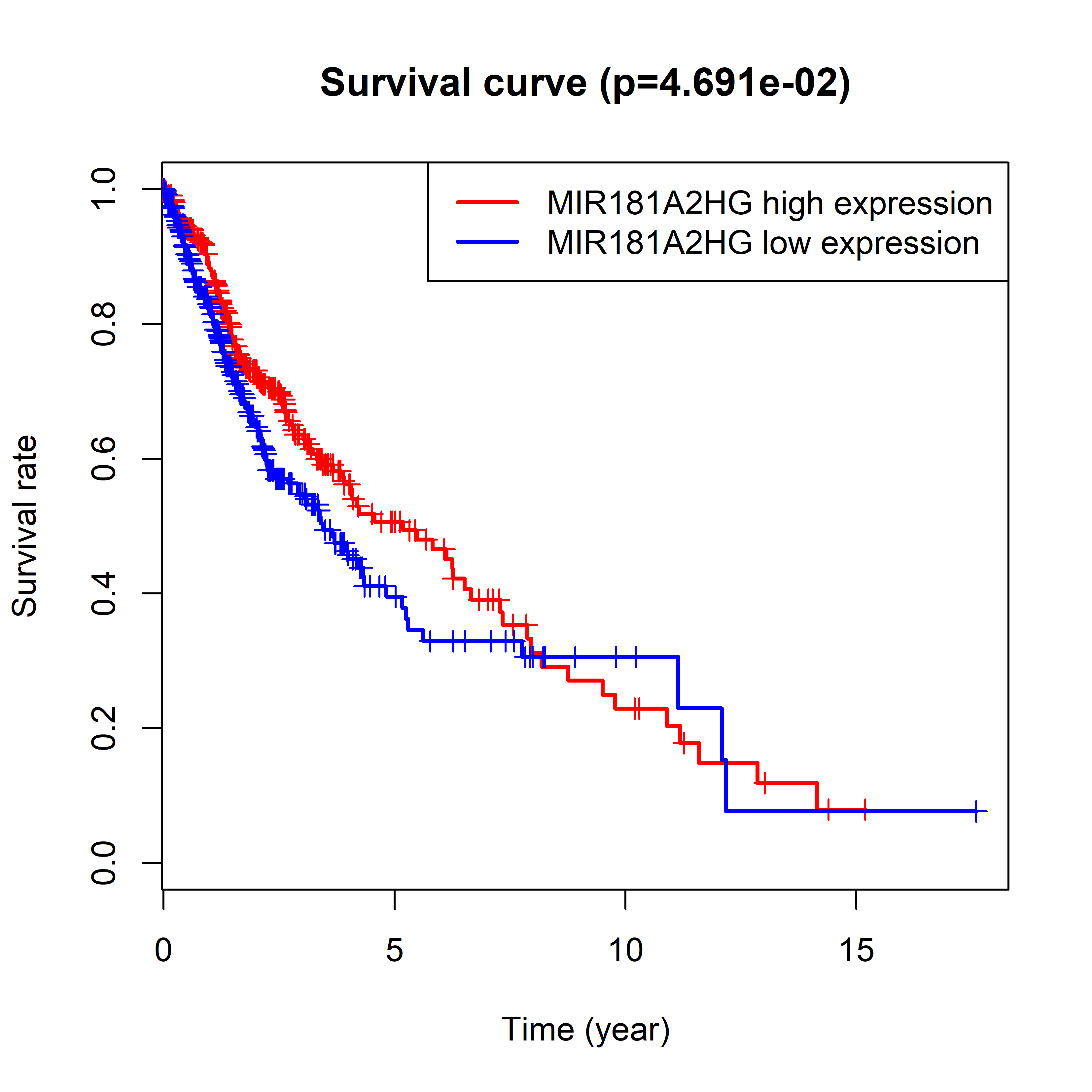

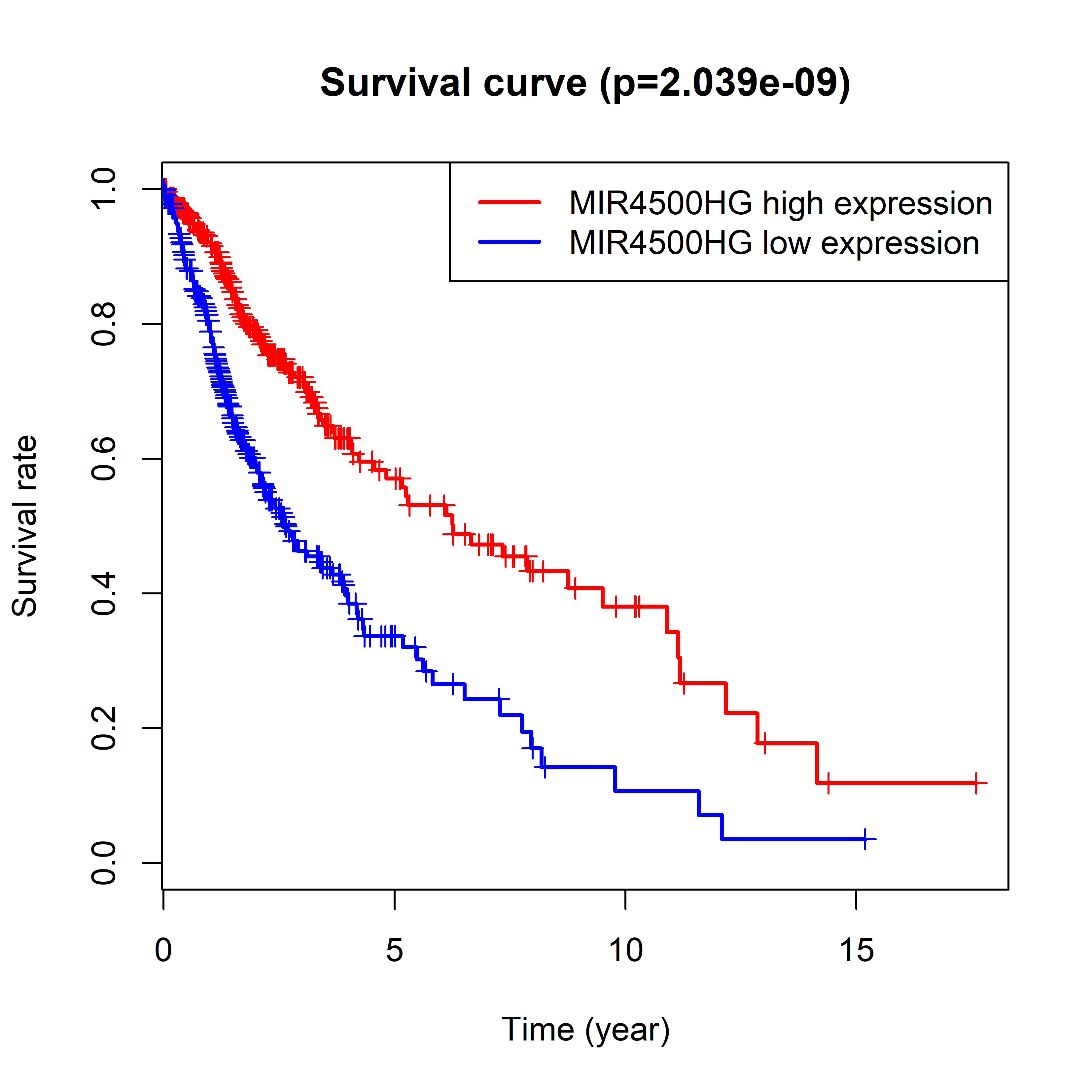

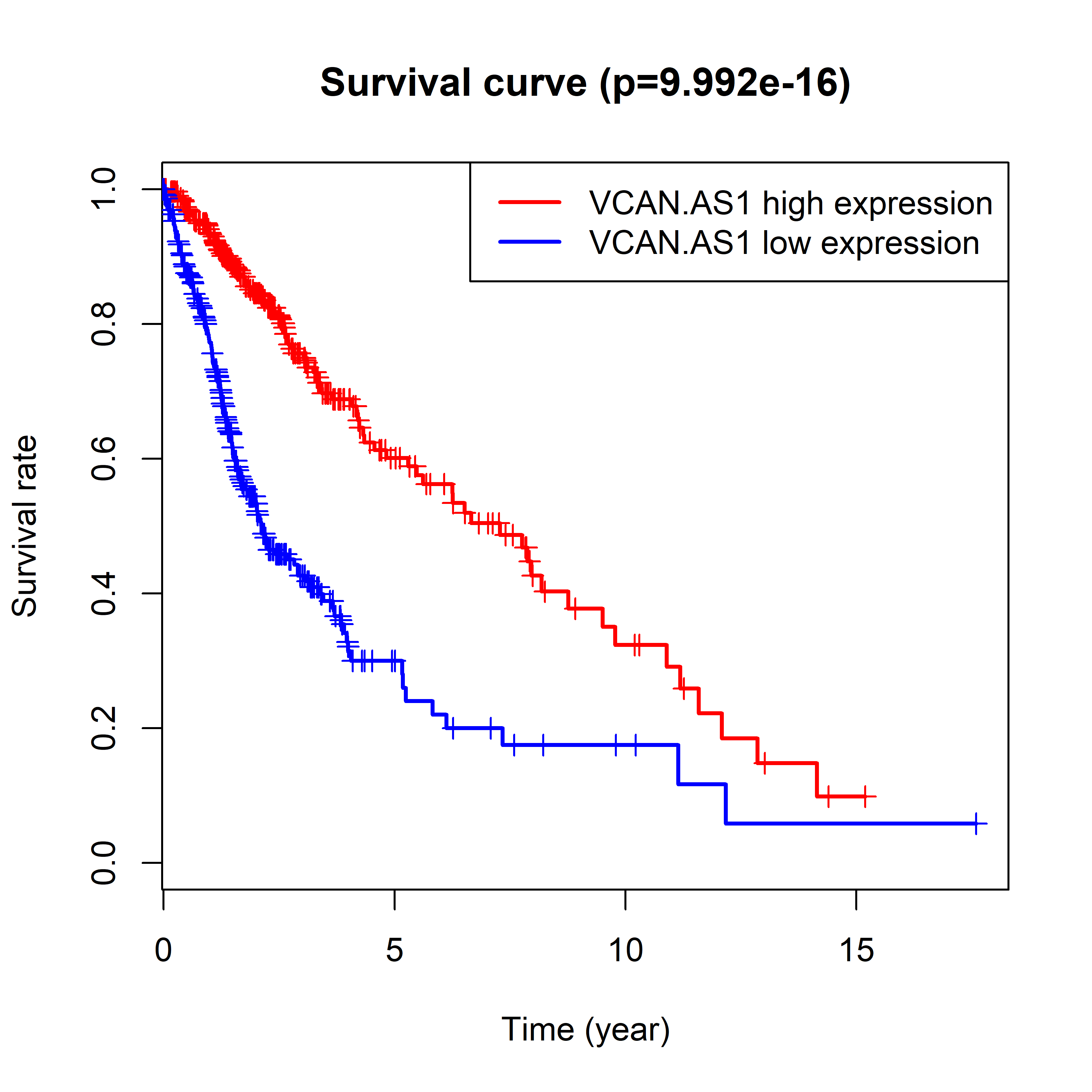

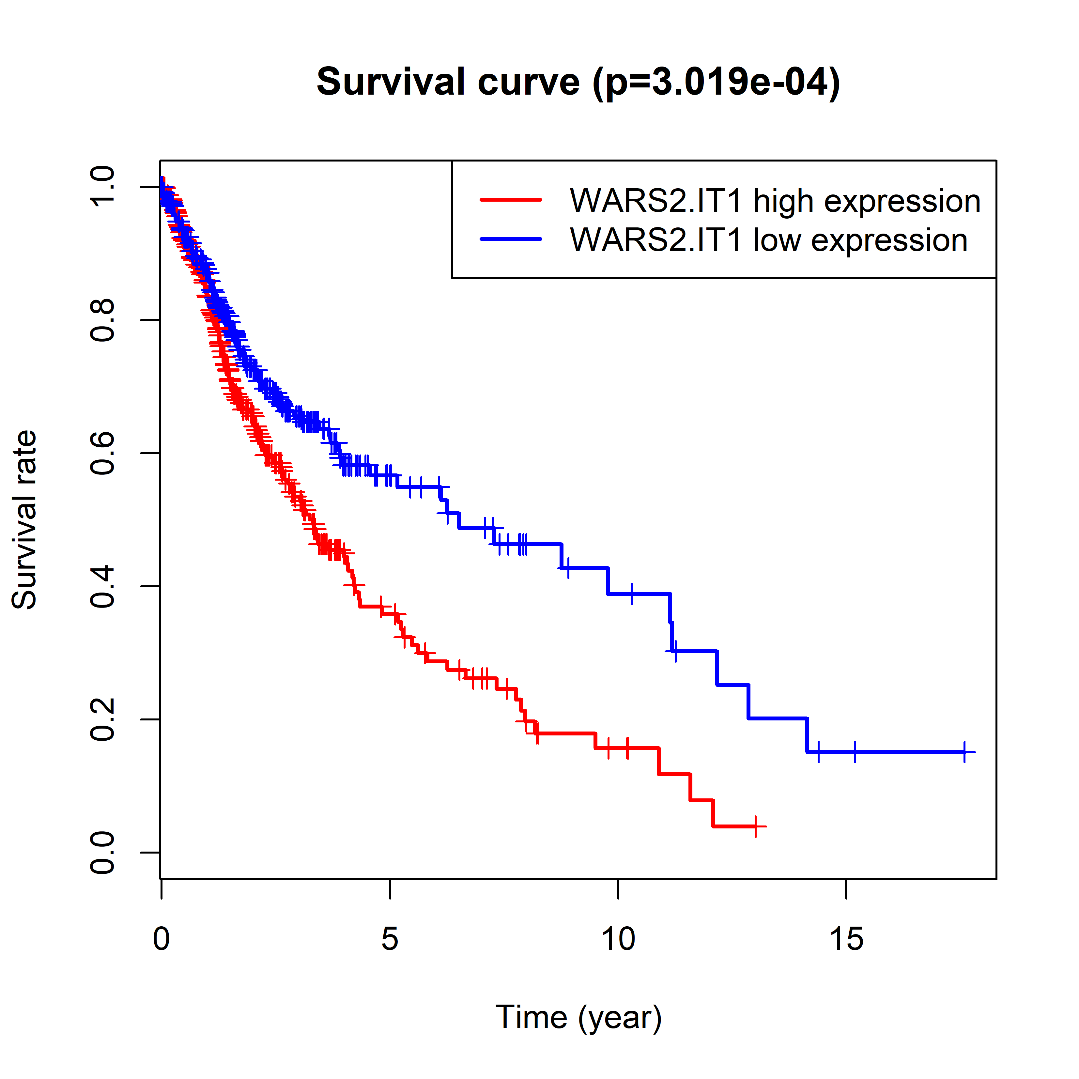
